# Inactivation of histone chaperone HIRA unmasks a link between normal embryonic development of melanoblasts and maintenance of adult melanocyte stem cells

**DOI:** 10.1101/2022.04.22.489166

**Authors:** Farah Jaber-Hijazi, Karthic Swaminathan, Kathryn Gilroy, Alexander T. Wenzel, Anthony Lagnado, Kristina Kirschner, Neil Robertson, Claire Reid, Neil Fullarton, Jeff Pawlikowski, Karen Blyth, Jill P. Mesirov, Taranjit Singh Rai, João F. Passos, Laura M. Machesky, Peter D. Adams

## Abstract

Histone chaperone HIRA is thought to play a role in both early development and aging, but little is known about connections between the two processes. Here, we explore this relationship using a lineage-specific knockout mouse model, *TyrCre::Hira^fl/fl^*, in which HIRA is deficient in the pigmentary system consisting of embryonic melanoblasts, postnatal melanocytes and melanocyte stem cells (McSCs). *Hira* knockout leads to reduced melanoblast numbers during embryogenesis, but wild type numbers of melanocytes at birth, normally functioning juvenile and young adult McSCs, and only a very mildly hypopigmented first hair coat. However, on closer analysis, *Hira* knockout melanocytic cells of newborn mice exhibit molecular markers characteristic of cell aging and proliferative deficits. As they age, *TyrCre::Hira^fl/fl^* mice display marked defects in McSC maintenance and premature hair graying. Importantly, these defects are only observed when HIRA is inactivated during embryogenesis, not post-natally. This genetic model illustrates how normal embryonic development lays the foundation for maintenance of adult tissue specific stem cells and so suppression of degenerative phenotypes of aging.

## Introduction

Epigenetic mechanisms play crucial roles in sustaining genome function and cell phenotype to drive development and promote healthy aging^1–3^. Histone variants, post-translational modifications on histone tails, nucleosome occupancy and other aspects of chromatin provide a flexible epigenome that helps cells respond and adapt to changing physiological and environmental conditions^4–8^.

HIRA is a histone chaperone, specifically targeted to the histone variant H3.3 which differs from H3.1 and H3.2 by only a few amino acids^9^. HIRA, which has significant homology to yeast Hir1p and Hir2p proteins, is one of several transcription units identified in the human 22q11 locus, the target of heterozygous deletions in human DiGeorge syndrome^10,11^. HIRA is a key member of the HIRA histone chaperone complex, also containing UBN1 and CABIN1, and cooperating with ASF1A^9,12,13^. At the molecular level, the HIRA histone chaperone complex deposits histone H3.3 into nucleosomes in a DNA replication independent manner, mainly at active promoters, enhancers and genic regions^9,14,15^. Nucleosomes carrying both H3.3 and H2A.Z histone variants are susceptible to disruption, marking active gene promoters^7^. HIRA and histone H3.3 are implicated in the cell response to DNA damage, both playing a role in DNA repair^16,17^. In addition, histone H3.3 has also been proposed to play a role in epigenetic memory of a transcriptionally activated state through serial rounds of cell division^18–20^.

HIRA is expressed during murine embryogenesis as early as embryonic stages E6.5-E7.5 during gastrulation^21,22^, and ubiquitous expression was observed at E8.5 with higher levels in the cranial neural folds^22^. Several studies indicate an essential role of HIRA and H3.3 in oogenesis and embryonic viability and development in mice^21,23–26^. H3.3 has a repressive role when trimethylated at lysine 27 at developmentally regulated bivalent domains in embryonic stem cells, suggesting an important role in differentiation^27^. Moreover, HIRA has been shown to be required for normal differentiation of various cell lineages, tissues and organs, such as neural cells, cardiomyocytes and the haematopoietic lineage^28–32^. HIRA and H3.3 have also been shown to play significant roles in cell and tissue aging. For example, HIRA is required for epigenetic integrity in cellular senescence, a stress-induced proliferation arrest that contributes to tissue aging^33,34^. H3.3 also accumulates in many tissues with age^35^, including neuronal and glial cells, orchestrating transcriptional programmes and maintaining physiological plasticity^5^. In sum, HIRA is a histone H3.3 chaperone involved in lineage differentiation, development and aging.

Accordingly, we set out to investigate the role of HIRA in the pigmentary system of mice, a very tractable model to study development and aging mechanisms. During embryogenesis, the precursors to melanocytes and melanocyte stem cells (McSCs) are highly proliferative unpigmented cells called melanoblasts. They begin specification from the neural crest at E9.5 and migrate dorsolaterally through the dermis between the somites and to populate the epidermis by E15.5^36,37^. Afterwards, melanoblasts colonise the developing hair follicles (HFs) where they differentiate into melanocytes and McSCs^38–40^.

In postnatal stages, HFs are in continuous turnover cycles of growth (anagen), regression (catagen) and rest (telogen)^39,41,42^. In all these phases the HF retains a permanent stem cell rich region, the bulge, but the anagen HF also has a lower region, the bulb. Hair colour is provided by melanin pigments produced from bulb melanocytes, which differentiate from non-pigmented McSCs residing in the bulge^39,42–44^. McSCs are usually quiescent in the telogen phase, but they initiate proliferation and differentiation in response to growth signals during the anagen phase^39,42,44^. Progressive loss of McSCs and therefore reduction in melanocyte numbers is one of the main causes of hair graying associated with aging^45^.

Besides the difference in their location or pigmentation, melanoblasts, McSCs and mature melanocytes can be distinguished from each other by gene expression patterns. In brief, before entering the growing HFs, melanoblasts express the microphthalmia-associated transcription factor (MITF), the master regulator of the melanocytic lineage, dopachrome tautomerase (DCT), paired box gene 3 (PAX3), tyrosinase (TYR), premelanosome protein (PMEL17), receptor tyrosine kinase (c-KIT) and SRY-box transcription factor (SOX10)^46,47^. After entering the HF, increasing expression of MITF and pigmentation genes, such as *Tyr* and *Pmell 7*, lead to differentiation into bulb melanocytes^38^, while diminishing levels of c-KIT and pigmentation genes lead to the formation of bulge McSCs^48,49^. Schwann cell precursors from the glial lineage have also been proposed as a source of melanoblasts, differentiating around E11.5 and migrating along the ventral pathway^50^. Phenotypes related to the differentiation, migration, proliferation, function, survival and aging of melanoblasts, melanocytes and McSCs are easy to observe, because defects often lead to a change in the patterns or levels of pigmentation^51,52^.

To probe the role of HIRA in the mouse pigmentary system, we generated a lineagespecific knockout mouse model, *TyrCre::Hira^fl/fl^*, in which McSCs, melanocytes and melanoblasts are deficient for HIRA. We find that HIRA is required during early and mid embryogenesis for proper development of melanoblasts. While defects caused by HIRA inactivation are normalized during late embryogenesis and are barely detectable in melanocytes and McSCs at birth and in young mice, they are manifest later as profound defects in McSC maintenance during adult aging. These results demonstrate a link between the integrity of embryonic development and phenotypes characteristic of healthy *vs* unhealthy aging.

## Results

### *Hira* knockout causes a reduction in melanoblast numbers during embryonic development

*Hira^fl/fl^* mice^34^ were crossed with *TyrCreA* (X-linked) or *TyrCreB* (autosomal) mice in which Cre recombinase is under the control of the *Tyr* gene promoter, allowing inactivation of *Hira* in the melanocytic lineage as early as embryonic days E9.5-10.5^53^. The generated *TyrCre::Hira^fl/fl^* mice were in turn crossed with *Rosa26-Lox-STOP-Lox (LSL)-tdTomato* mice to track Cre recombinase activity^54^. Recombination efficiency was confirmed in E15.5 embryos by co-localization of DCT, a melanoblast marker, and tdTomato in the epidermis (**Fig. 1a**). Not all tdTomato+ cells are DCT+ because, as reported previously^53^, the *Tyr* promoter also directs expression in some other non-melanocytic lineages (**Fig. 1a; also see Figure 2**). These results confirm *TyrCre*-directed recombination in embryonal DCT+ cells.

**Fig. 1.**
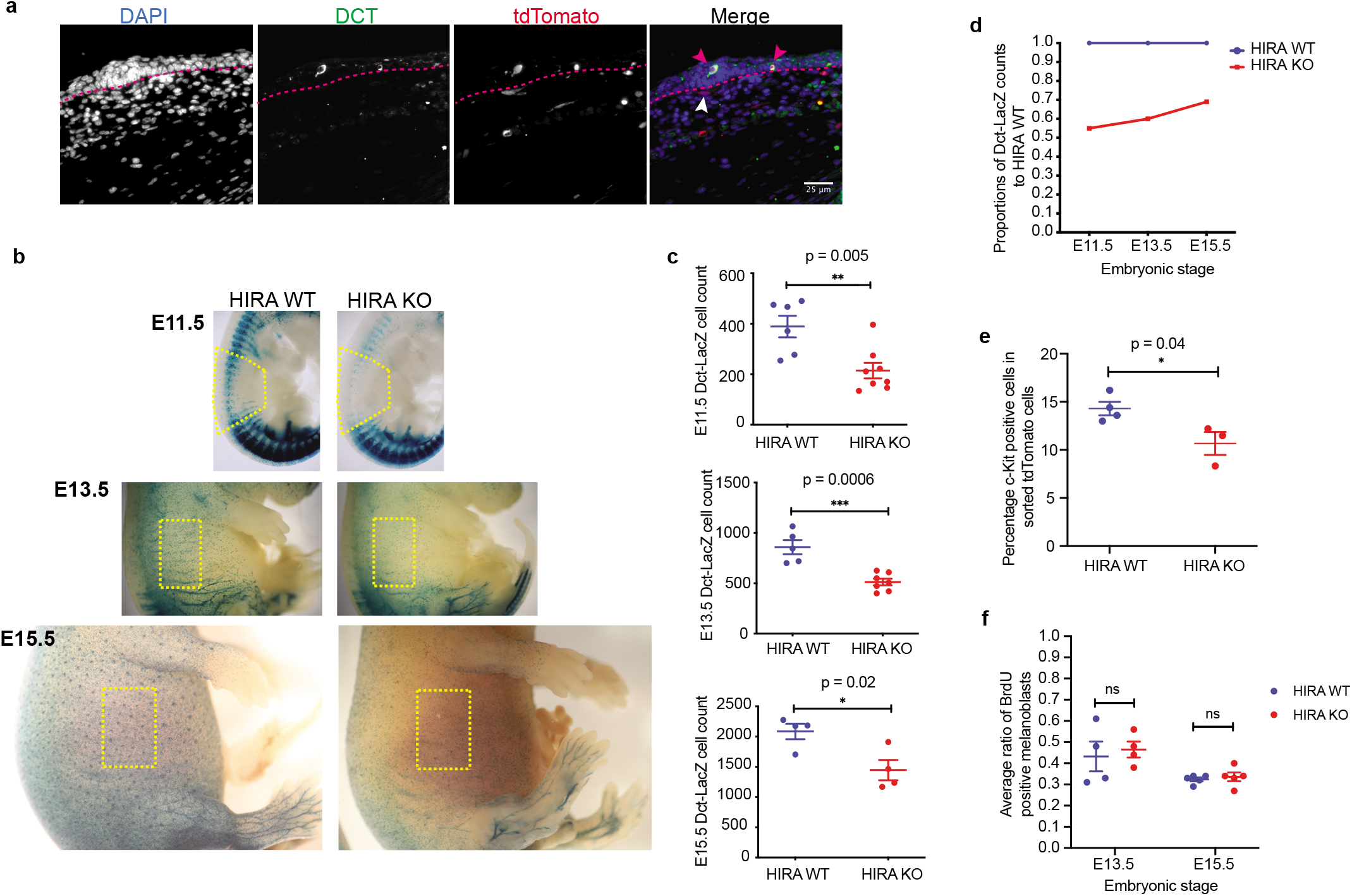
*Hira* knockout causes a reduction in melanoblast numbers during embryonic development. **a,** Immunofluorescence in *TyrCre::Hira^wt/wt^* (HIRA WT) E15.5 mouse embryos showing tdTomato expression, identified by RFP antibody (red), specifically in melanoblasts (magenta arrow heads) in the epidermis identified by DCT stain (green). The epidermis and dermis of the skin are separated with dotted magenta line, epidermis on top. White arrowheads point to non-melanoblast dermal tdTomato+ cells. Antibody specificity was confirmed with Cre negative or tdTomato negative samples as negative controls. Scale bar: 25 μm. **b,** X-gal stain in E11.5, E13.5 and E15.5 in *TyrCre::Hira^fl/fl^* (HIRA WT) and *TyrCre::Hira^fl/fl^* (HIRA KO) *Dct::LacZ* embyos showing melanoblast distribution. Embryos are shown in real relative size. Yellow boxes represent the areas quantified. **c,** Quantification of yellow boxes in (**b**). Error bars represent the standard error of the mean (s.e.m). Embryo numbers at E11.5, WT n = 6 and KO n = 8; E13.5, WT n = 5 and KO n = 7; E15.5, WT n = 4 and KO n = 4 from at least 2 litters for each timepoint. **d,** Quantification of Dct-LacZ counts (**c**) in HIRA KO and HIRA WT normalized to WT at each time point, showing a diminishing difference in the number of HIRA KO to HIRA WT melanoblasts as gestation advances. **e,** Percentage of melanoblasts that are c-KIT positive, selected using a BV711-conjugated antibody to c-KIT, within the tdTomato populations in *tdTomato* HIRA WT and HIRA KO E15.5 embryos. Melanoblasts were quantified by FACS profiling of single trunk skin cell suspensions. *tdTomato* HIRA WT, n = 4; *tdTomato* HIRA KO, n = 3. See **Supplementary** Fig. 1 for the gating strategy used for quantification. **f,** Percentage of BrdU positive melanoblasts quantified by BrdU and DCT antibody stains in E13.5 and E15.5 embryo cross sections following a 2 hour BrdU pulse, with n = 4 for each genotype at E13.5 (p = 0.7) and n = 5 for each genotype at E15.5 (p=0.6). Scatter dot plot data in **c, e** and **f** were analysed using an unpaired *t*-test showing mean ± s.e.m. ns: non-significant.

To investigate a role for HIRA in DCT+ embryonic melanoblasts, we crossed *TyrCre::Hira*^fl/fl^ mice with *Dct::LacZ* reporter mice^55^ in order to track melanoblasts through embryonic development. We compared melanoblast number and distribution between *TyrCre::Hira^fl/fl^ Dct::LacZ* (HIRA KO) and control *TyrCre::Hira^wt/wt^ Dct::LacZ* (HIRA WT) at E11.5, E13.5 and E15.5 of embryogenesis, by staining with X-gal. This showed that HIRA KO embryos have a significant reduction in the number of melanoblasts compared to HIRA WT embryos at all three stages (**Fig. 1b, c**). Interestingly, the ratio of melanoblast numbers in HIRA KO to that in HIRA WT was 0.55 at E11.5, 0.59 at E13.5 and 0.69 at E15.5, suggesting a greater effect of HIRA KO at earlier stages of embryogenesis (**Fig. 1c, d**). We also quantified melanoblast numbers by flow cytometry (**Supplementary Fig. 1**). To identify melanoblasts, we stained cell suspensions from individual *LSL-tdTomato* HIRA KO and WT E15.5 embryo skins **(Fig. 1a)** with a BV711-conjugated antibody against c-KIT, widely used as a melanoblast cell-surface marker^56–58^. We found that the percentage of c-KIT+ tdTomato+ cells was significantly reduced in HIRA KO samples (10.7% ±2%, n = 3) compared to HIRA WT samples (14.3% ±1.4%, n = 4) (**Fig. 1e**), indicating a reduction in melanoblast numbers, consistent with that observed in the *Dct-LacZ* reporter system.

To determine whether HIRA had an effect on melanoblast proliferation that might account for their decreased number in HIRA KO embryos, E13.5 and E15.5 embryos were harvested after a 2-hour BrdU pulse to pregnant dams. Cross sections were stained with anti-BrdU and anti-DCT antibodies to determine the percentage of proliferating melanoblasts in both the upper and lower halves of the body along the axial plane. The average percentage of BrdU positive melanoblasts was not significantly different between HIRA KO and HIRA WT embryos; (46.5% ±7.7% (n = 4) in HIRA KO and 43.4% ±14.2% (n = 4) in HIRA WT embryos at E13.5; 33.7% ±4.7% (n = 5) in HIRA KO and 32.5% ±2.1% (n = 5) in HIRA WT embryos at E15.5) (**Fig. 1f**). In sum, at E13.5-15.5, embryonic HIRA KO melanoblasts show similar proliferation to WT melanoblasts, but reduced total number.

### HIRA is required for melanoblast development

To better understand the defect in melanoblast development in HIRA KO embryos, we isolated tdTomato+ cells from trunk skin of *TyrCre::Hira^wt/wt^:tdTomato* (WT) and *TyrCre::Hira^fl/fl^:tdTomato* (KO) E15.5 embryos (3 embryo replicates each genotype). The purified cells were 95% tdTomato+ (**Fig. 2a** and **Supplementary Fig. 2a**). Transcriptomes of WT and KO melanoblasts were compared by single cell RNA sequencing (scRNAseq) on the 10x Chromium platform (see **Supplementary Table 1** for details on the samples). Data were analyzed by 10x Cell Ranger, cLoupe and Seurat^59^. Comparing data from the individual WT and KO replicates to the t-SNE plot generated from all pooled WT and KO samples showed that the cell sub-clusters in WT and KO samples were broadly similar (**Fig. 2b**).

**Fig. 2.**
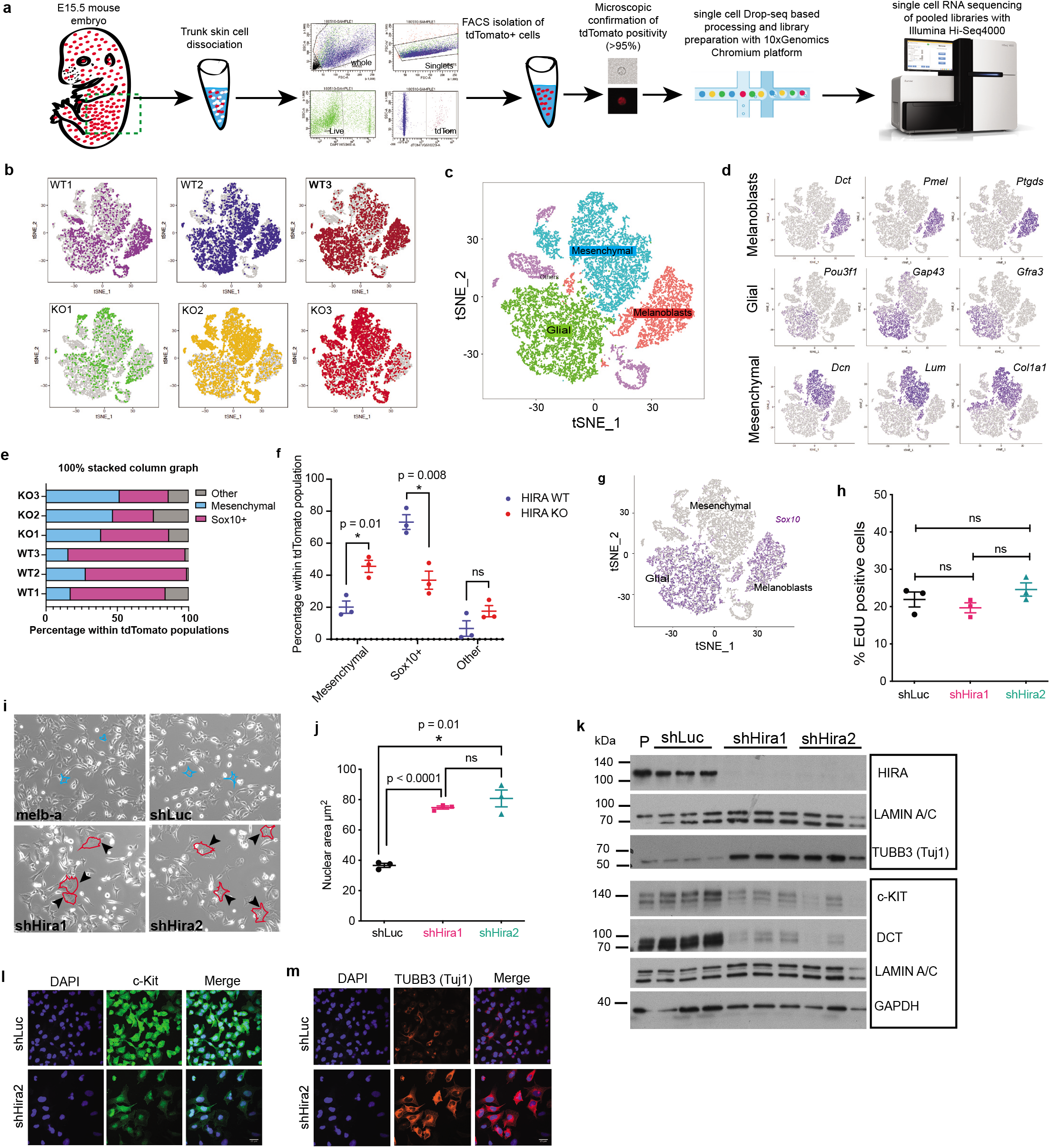
HIRA is required for melanoblast development. **a,** Scheme of single cell RNA sequencing method. 3 *TyrCre::Hira^wt/wt^:tdTomato* (WT) and 3 *TyrCre::Hira^fl/fl^:tdTomato* (KO) embryos were individually processed. **b,** t-SNE plots showing clusters based on gene expression among a total of 8655 single cells from 3 WT and 9743 single cells from 3 KO embryos. Cells from all 3 WT and 3 KO replicates are superimposed in each plot with the cells of each sample highlighted in a specific colour on each of the 6 plots. **c,** t-SNE plot of all 6 pooled samples showing 3 major cell types. **d,** t-SNE plots showing the expression of representative marker genes in the 3 major cell types. Expression level is color-coded in purple. **e,** 100% stacked column graph displaying the percentage of *Sox10+* and mesenchymal cells within the tdTomato population in each of the 6 samples. Cell counts and percentages can be found in **Supplementary Table 5. f,** Scatter dot plots showing the percentages of cells in each of the populations in **e** in WT *versus* KO samples. **g,** t-SNE plot displaying *Sox10* expression level in melanoblasts and glial cells **h,** Scatter dot plots displaying the average percentage of EdU positive nuclei in melb-a cells 10 days after infection with shHira1 and shHira2 lentiviral vectors, with shLuc as control. 3 independent replicates each (3 culture plates each each infected separately). **i,** Brightfield images of representative melb-a cells one week following infection as in (**h**). Uninfected (non-drug selected) parental cells are labelled as melb-a. Black arrows indicate enlarged cells (also traced in red) compared to parental and shLuc control (some cells traced in blue for comparison). Magnification 10x. **j,** Scatter dot plots representing the average nuclear area of cells in (**i**) as measured with ImageJ. **k,** Representative Western blots of HIRA, TUBB3 and the melanoblast-specific proteins c-KIT and DCT in lysates from shLuc, shHira1 and shHira2 melb-a cells, 3 independent replicates each, 10 days following infection and drug selection. GAPDH and Lamin A/C were used as loading and sample integrity controls, respectively. Blots performed in the same gel or a re-probed membrane are boxed together. Molecular weights were marked using Spectra multicolour broad range protein ladder. P: parental. **l,** c-KIT immunofluorescence (green) melb-a cells 10 days post-infection. **m,** TUBB3 immunofluorescence on same cells in (**l**). Scale bars: 25 μm. Scatter dot plot data in **f**, **h** and **j** were analysed using an unpaired *t-*test showing mean ± s.e.m. ns: non-significant (p>0.05).

Based on expression of sub-cluster-specific marker genes and biological or functional pathways such as the cell cycle, 13 sub-clusters (numbered 0-12) were observed in the pooled WT and KO cells (**Supplementary Fig. 2b, c**). However, when only characteristic lineagespecific genes were considered, we identified three main populations of cells resembling melanoblasts, glial cells and mesenchymal cells (**Fig. 2c, d**). These 3 major subclusters were distinguished by distinct gene expression profiles: For example, melanoblasts expressed *Dct, Pmel* and *Ptgds;* glial cells *Pou3f1, Gap43, Gfra3;* and mesenchymal cells *Dcn, Lum, Col1a1* (**Fig. 2d)**. More examples of cluster-specific distinguishing genes were generated from WT populations only and can be found in **Supplementary Tables 2-4** and **Supplementary Fig. 2d-f.** In WT embryos, a minority of tdTomato+ cells were mesenchymal (**Fig. 2e, f and Supplementary Table 5**), consistent with previous reports that some mesenchymal cells are derived from the neural crest^60^. Since scRNAseq shows that these cells do not express *Tyr* at E15.5 (**Supplementary Fig. 2d**), presumably these cells are derived from a neural crest lineage that previously expressed *Tyr* to irreversibly activate expression of *tdTomato*. More specifically, Schwann cell precursors of the glial lineage, also previously expressing *Tyr,* may transdifferentiate into mesenchymal cells as well as melanoblasts^50,61^. In WT embryos, the majority of tdTomato+ cells were *Sox10+* melanoblasts and glial cells (74.7%; **Fig. 2e-g and Supplementary Table 5**). This is consistent with a neural crest-derived SOX10+ progenitor as a precursor to both melanoblasts and glial cells^62^, and the previously demonstrated activity of the *Tyr* promoter in these cell types^53,63,64^.

Although tdTomato+ cells from WT and KO embryos displayed broadly similar cell sub-clusters, we observed a change in proportions from WT to KO (**Fig. 2e-g** and **Supplementary Table 5**). Mean melanoblast percentage was significantly reduced from 30.3% ±6.2% in HIRA WT to 12.5% ±4.6% in HIRA KO, in line with our previous observations by flow cytometry analysis and the *Dct-LacZ* reporter (**Fig. 1b-e)**. When we assessed combined *Sox10+* cells (melanoblasts and glial cells), we observed a significant reduction in the mean *Sox10+* population from 73.22% ±8% in HIRA WT to 36.9% ±9.8% in HIRA KO samples (**Fig. 2e, f**). These data are also in agreement with the downregulation in expression of various other melanoblast and glial-specific genes, such as *Dct, Ptgds, Fabp7* and *Gfra3, Mpz* and *Mbp* as observed in differential expression (DE) analysis (**Supplementary Tables 6-8**). The decrease in *Sox10+* cells in KO populations was accompanied by a significant increase in the percentage of mesenchymal cells (20.1% ±6.6% in HIRA WT to 45.6% ±6.6% in HIRA KO (**Fig. 2e, f, Supplementary Fig. 2g, h and Supplementary Table 5**). This was accompanied by an upregulation of the mesenchymal genes *Col1a1, Col3a1, Lum* and *Dcn* as observed from DE analysis of KO against WT melanoblast clusters (**Supplementary Table 6**). Moreover, the KO mesenchymal population had an increased expression of epithelial-to-mesenchymal transition (EMT) genes (**Supplementary Fig. 2i**), indicating that WT and KO cells were qualitatively distinct. Therefore, loss of HIRA not only decreases the proportion of tdTomato+ cells with a melanoblast identity, but the *Sox10+* lineage as a whole, accompanied by a corresponding increase in atypical mesenchymal cells

To confirm these results in a complementary model *in vitro*, we examined the consequence of *Hira* knockdown in mouse melb-a melanoblasts^65^. For this, we used two different lentivirus-encoded shRNAs to knockdown *Hira*, shHira1 and shHira2, and analysis was performed 8-10 days following infection and drug selection. shRNA targeted to luciferase (shLuc) was used as a negative control. Knock down of expression of HIRA protein was confirmed by western blot, and this was accompanied by downregulation of another member of the HIRA chaperone complex, CABIN1, as previously reported^12^ (**Supplementary Fig. 2j**). Consistent with data from mouse embryos (**Fig. 1f**), knockdown of *Hira* had no significant effect on DNA synthesis, analysed following a 2-hour EdU pulse (**Fig. 2h**). However, when compared to uninfected (parental) and shLuc melb-a cells, some shHira1 and shHira2 cells appeared to have a more fibroblastic/mesenchymal morphology, consistent with the scRNAseq data (**Fig. 2i, j**). In line with the scRNAseq, we observed a decrease in DCT and SOX10 expression in shHira1 and shHira2 cells (**Fig. 2k** and **Supplementary Fig. 2k)**. Moreover, we found a striking reduction in the expression of other melanoblast/cyte-specific markers c-KIT and MITF (**Fig. 2k, l and Supplementary Fig. 2l**). On the other hand, we found an increase in the expression of the TUBB3 protein (TUJ1) (**Fig. 2k, m**), but no effect on other tubulin isoforms (**Supplementary Fig. 2m**). Expression of TUBB3 has been previously associated with neural, melanocytic and EMT fates^31,66–68^. Increased expression of the *Tubb3* mRNA was also observed in scRNAseq DE analysis both in melanoblast and glial KO vs WT clusters (**Supplementary Tables 6-8**). In conclusion, all our data show that HIRA is required for the proper development of the melanoblast lineage and its inactivation leads to an aberrant differentiation programme.

### Knockout of *Hira* does not affect the distribution and number of melanocytes and McSC function in newborn mice

In light of this clear impediment to proper development of HIRA-deficient melanoblasts, we next set out to understand the consequences for McSC and melanocytes in newborn mice and young and aging adults. Surprisingly, given impaired development of the melanoblast lineage during early embryogenesis, *TyrCre::Hira^fl/fl^* (HIRA KO) and *TyrCre::Hira^wt/wt^* (HIRA WT) pups were almost indistinguishable in colour of their first hairs, both in *TyrCreX* and *TyrCreB* models (**Fig. 3a**), indicating the presence of functional melanocytes at this stage. To confirm *TyrCre*-directed recombination efficiency in newborn mice, we used the tdTomato reporter system and found that DCT+ melanocytes in anagen hair follicles of postnatal day 10 (P10) pups uniformly and specifically expressed tdTomato (**Fig. 3b**). In addition, as a functional readout for HIRA inactivation, we used immunofluorescence to assess the nuclear levels of H3.3. Interestingly, in HIRA WT mice, we observed marked enrichment of H3.3 specifically in DCT+ melanocytes and anatomical dermal papillae of hair bulbs, compared to other cells (**Fig. 3c**). Both of these cell types are non-proliferative^69,70^, consistent with DNA replication-independent deposition of H3.3. However, enrichment in melanocytes was absent in HIRA KO melanocytes (**Fig. 3c**), but retained in dermal papilla, consistent with lineage-specific HIRA inactivation in melanocytes and confirming a requirement for HIRA for deposition of H3.3 in these cells. Consistent with near-normal hair coat color of HIRA KO mice (**Fig. 3a**), immunostaining of melanocytic cells using anti-DCT antibody confirmed a similar distribution and number of DCT+ cells between HIRA WT and HIRA KO in P1 and P10 pups (**Fig. 3d**). Quantitation confirmed no significant difference in the mean number of melanocytes per anagen hair follicle in P10 HIRA WT (6.24 ±1 DCT+ cells, n = 3 pups) and HIRA KO (6.97 ±1.6 DCT+ cells, n = 3 pups) (**Fig. 3e**). We made similar observations in P1 pup skin using MITF and SOX10 immunostaining in consecutive sections to DCT stains (**Fig. 3f**). Quantification of DCT+ cells in P1 pup skin, a mixture of melanoblasts in the epidermis and McSCs and differentiating melanocytes in newly forming hair follicles, also showed that their numbers per unit area (x10^-4^) were not significantly different between HIRA WT (3.2 ±0.14, n = 3) and HIRA KO (2.9 ±0.25, n = 3) skins (**Fig. 3g**).

**Fig. 3.**
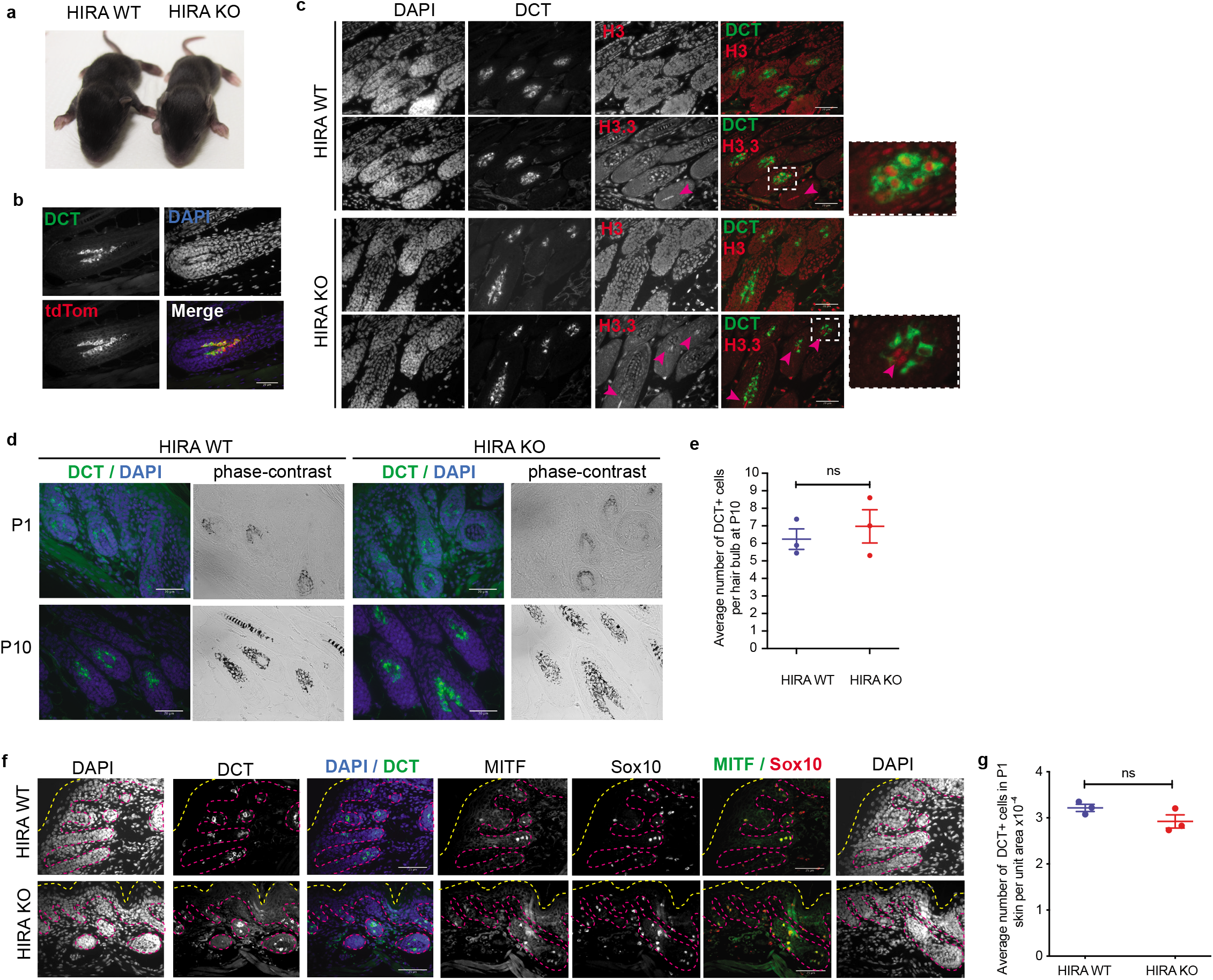
Knockout of *Hira* does not affect the distribution and number of melanocytes and McSC function in newborn and young adult mice. **a,** Images of *TyrCre::Hira^wt/wt^* (HIRA WT) and *TyrCre::Hira^fl/fl^* (HIRA KO) P11 pups showing coat colour of first hairs. **b,** Efficient recombination in HIRA KO mice as shown by overlapping expression of tdTomato (red), induced by Cre-recombinase, with DCT+ hair bulb melanocytes (green). Skin sections display hair at the anagen phase of pups at postnatal day 10 (P10). Scale bar: 20 μm. **c,** Distinct expression of H3.3 (red) in specific cell types, including melanocytes (green), and dermal papillae (magenta arrow heads), in HIRA WT skin, as opposed to Histone H3 (red) which is more homogeneous in all skin cells. H3.3 staining in melanocytes is lost upon HIRA KO, but not in dermal papillae, while histone H3 expression remains unaffected. Skin longitudinal sections from P10 pups. Scale bar: 20 μm. **d,** DCT (green)/DAPI (blue) immunohistochemistry and corresponding phase contrast images showing pigmented melanocytes in growing hair bulbs of HIRA WT and HIRA KO mouse skin at P1 and P10. Scale bars 20 μm. **e,** Scatter dot plot showing the average number of melanocytes, measured by DCT positivity, in the hair bulbs from P10 HIRA WT and HIRA pup skin. n = 3 pups per genotype; p = 0.7. An average of 11-12 hair bulbs were counted pair sample. **f,** Representative image of melanocytic distribution in HIRA WT and HIRA KO P1 skin using 3 melanocytic markers: DCT, MITF and SOX10. DCT/DAPI and MITF/SOX10/DAPI images are taken from immunofluorescence done on consecutive paraffin sections. Outer layer of the epidermis is traced by a dotted yellow line. Basement membrane and hair follicles are traced by a dotted magenta line. Scale bar: 25 μm. **g,** Quantification of DCT positive cells, displayed in (**f**), per unit area (pixel^2^) of a skin field of view at 20x magnification. n = 3 samples per genotype; p = 0.1. Scatter dot plot data in **e** and **g** were analysed using an unpaired *t-*test showing mean with ± s.e.m. ns: non-significant.

HIRA KO mice, when compared to HIRA WT mice, maintained their coat colour and DCT+ cell number and distribution, with only a mild hypopigmentation, following the first real anagen at 4 weeks of age (**Supplementary Fig. 3a,b**). This indicates that, at least over the first few hair cycles, HIRA KO McSCs of young mice are capable of regenerating pigment-generating melanocytes. In conclusion, the knockout of *Hira* in the melanocytic system has a mild effect, if any, on melanocyte number and distribution and McSC function in newborn mice and young adults.

### Increased TAF in *Hira* knockout melanocytes and loss of function under stressful conditions

So far our results showed that despite having reduced melanoblast numbers in earlier developmental stages, newborn and young adult HIRA KO mice have seemingly normal functioning McSCs and similar numbers of melanocytic cells as HIRA WT mice. To better compare the phenotype and function of newborn WT and HIRA KO melanocytes, we isolated these cells from P1-P3 pups and cultured them *ex vivo*, as previously described in a medium containing the mitogen tetradecanoylphorbol-12-acetate (TPA)^71^. While HIRA WT melanocytes, identified by pigment and DCT expression, showed steady growth over 3 weeks in culture, HIRA KO melanocytes consistently failed to expand in number (**Fig. 4a**). Moreover, after a 48 hour pulse of EdU following 1, 2 or 3 weeks in culture, fewer HIRA KO melanocytes incorporated EdU compared to HIRA WT, consistent with their inability to expand in number (**Fig. 4a, b**). This indicates a melanocyte functional defect in HIRA KO not previously observed *in vivo*, and only revealed when the cells are challenged to proliferate under *ex vivo* conditions. Significantly, conditional inactivation of *Hira in vitro* by addition of 4-hydroxytamoxifen (4-OHT) to melanocytes isolated from *TyrCreERT2::Hira^fl/fl^* mice had no observable effect, suggesting that the *in vitro* defect observed in constitutive HIRA KO *TyrCre::Hira^fl/fl^* melanocytes necessarily originates during embryogenesis (**Supplementary Fig. 4a-d**).

**Fig. 4.**
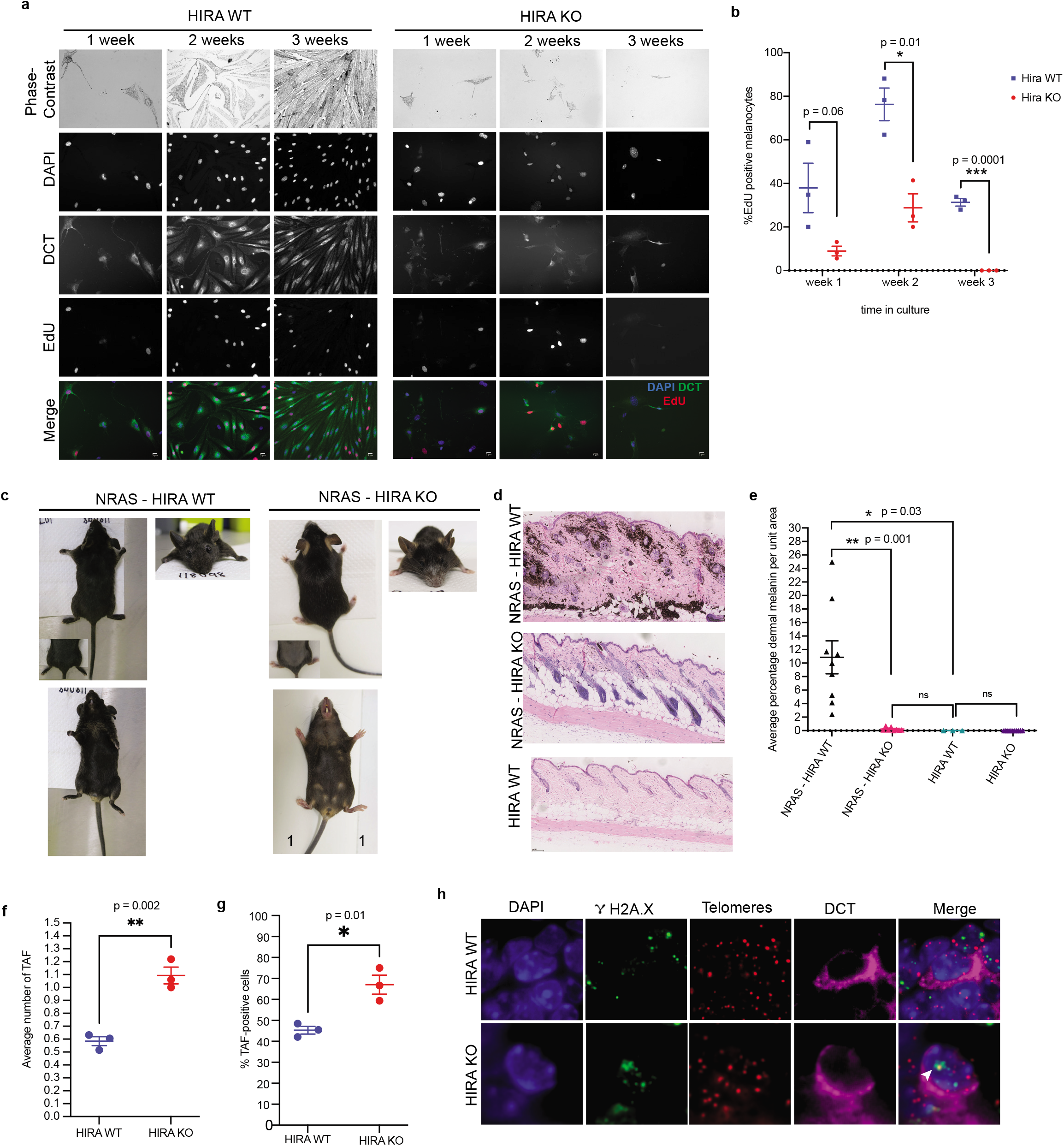
Increased TAF in HIRA knockout melanocytes and failure to thrive under stressful conditions. **a,** Comparison of *ex vivo* growth of primary melanocytes cultured from back skin of *TyrCre::Hira^wt/wt^* (HIRA WT) and *TyrCre::Hira^fl/fl^* (HIRA KO) P3 pups at 1, 2 and 3 weeks following seeding. Melanocytes were identified both by pigment and DCT positivity (green). EdU positive nuclei (red) were used to quantify proliferation. Scale bar: 5 μm. **b,** Scatter dot plots showing the percentage of EdU positive nuclei. Proliferation was assessed by measuring the percentage of EdU positive nuclei in melanocytes after a 48 hour EdU pulse. Cells were fixed at 1 week, 2 weeks or 3 weeks post culture. Cells were not passaged during the study period. n = 3 independent biological replicates per genotype cultured from different pups. **c,** Images of 5 month old *TyrNras::TyrCre::Hira^wt/wt^* (NRAS-HIRA WT) and *TyrNras::TyrCre::Hira^fl/fl^* (NRAS-HIRA KO) mice displaying differences in the colour of hair and skin of the snout, ears, chin, back, belly, feet and tail. **d,** H&E of longitudinal skin sections displaying difference in pigmentation between NRAS-HIRA WT, NRAS-HIRA KO and HIRA WT skins. **e,** Scatter dot plots showing the percentage of dermal melanin per unit area of skin. Quantification of skin melanisation in the dermis was measured by calculating the percentage area covered in brown pigment using an ImageJ plugin designed for this purpose. Only dermal region was quantified; hair follicles, fatty regions epidermis were excluded. Number of mice used for each genotype: NRAS-HIRA WT, n = 9; NRAS-HIRA KO, n = 8; HIRA WT, n = 3; HIRA KO, n = 10. **f,** Average number of telomere-associated DNA damage foci (TAF) in DCT positive cells (21 to 35 cells per sample) in P1 pup skins from 3 HIRA WT and 3 HIRA KO samples. **g,** Percentage of TAF positive melanocytes per genotype. **h,** Representative images of *γ*H2A.X immuno-FISH (Green, *γ*H2A.X; red, telomeres) in the nuclei of melanocytes (magenta). White arrowhead: colocalisation of *γ*H2A.X and telomeres indicating TAF. Scatter dot plot data in **b, e, f** and **g** were analysed using an unpaired *t-*test showing mean ± s.e.m. ns: non-significant (p > 0.05)

In order to similarly challenge proliferative capacity of the melanocytic cells *in vivo*, we used the *TyrNras* mouse model in which *Tyr*-driven expression of an *Nras^Q61K^* oncogene leads to hyperpigmentation of the skin due to excessive post-natal proliferation of bulb melanocytes and their expansion into dermal regions of the skin^72,73^. While the hyperpigmentation phenotype was observed as expected in *TyrNras::TyrCre::Hira^wt/wt^* (NRAS-HIRAWT) mice, it was clearly suppressed in *TyrNras::TyrCre::Hira^fl/fl^* (NRAS-HIRA KO) mice, as observed particularly in the snout, chin, ears, tail, feet and shaved skin (**Fig. 4c**). Haematoxylin and eosin (H&E) stained longitudinal skin sections also showed that the dermal pigmentation observed in NRAS-HIRA WT was not observed in NRAS-HIRA KO mice (**Fig. 4d**), and indeed in the latter dermal pigmentation was substantially and significantly reduced to levels similar to HIRA WT mice (**Fig 4d, e and Supplementary Fig. 4e**). These observations in adult mice were also evident in pups as young as P10 as shown by DCT staining (**Supplementary Fig. 4f**), confirming that decreased pigmentation in HIRA KO reflects decreased number of melanocytes.

We reasoned that this impaired response of newborn HIRA KO melanocytes to proliferative challenge both *in vitro* and *in vivo* might reflect elevated molecular stress in these cells. Specifically, telomeric dysfunction is known to limit the proliferative capacity of cells. Therefore, we examined a marker of telomere dysfunction, telomere-associated DNA damage foci (TAF)^74^, by measuring the number of *γ*H2A.X foci at telomeres in DCT+ cells of P1 pups. We found that the mean number of TAFs per DCT+ cell was 0.58 ±0.06 in HIRA WT, but was significantly higher in HIRA KO cells (1.09 ±0.11) (**Fig. 4f**). Moreover, the percentage of TAF+ DCT+ cells was also significantly higher in HIRA KO pups (67.01% ±7.82%) than HIRA WT pups (45.35% ±3.2%) (**Fig. 4g, h**). Together, these data indicate that newborn HIRA-deficient McSCs and melanocytes exhibit features of molecular stress and impaired proliferation potential *in vitro* and *in vivo.*

### HIRA is required during embryonic development for melanocyte stem cell maintenance in adults and suppression of hair greying during aging

Prolonged McSC function and cyclical proliferation is necessary for maintenance of McSCs and regeneration of mature differentiated melanocytes through each hair cycle, and hence suppression of hair greying over the lifecourse. Based on our results, we hypothesized that HIRA-deficient McSCs would be unable to perform these functions over the long term, despite their apparent normal function over the first few hair cycles **(Fig. 3)**. Indeed, consistent with this idea, by 20 months old, naturally aged HIRA KO mice showed a strong premature hair graying phenotype with pigment loss most notable in the snout, belly and lower parts of the back (**Fig. 5a**). Importantly, in line with the previous *in vitro* experiments, this premature hair graying phenotype was not observed even 18 months after *Hira* inactivation by 4-OHT administration to 6 week old conditional KO *TyrCreERT2::Hira^fl/fl^* mice (**Supplementary Fig. 5a**). In this case, recombination was confirmed, both in McSCs and melanocytes, by tdTomato expression. (**Supplementary Fig. 5b, c**). This again indicates that the pigmentation defects in KO *TyrCre::Hira^fl/fl^* mice necessarily originate during embryonic development.

**Fig. 5.**
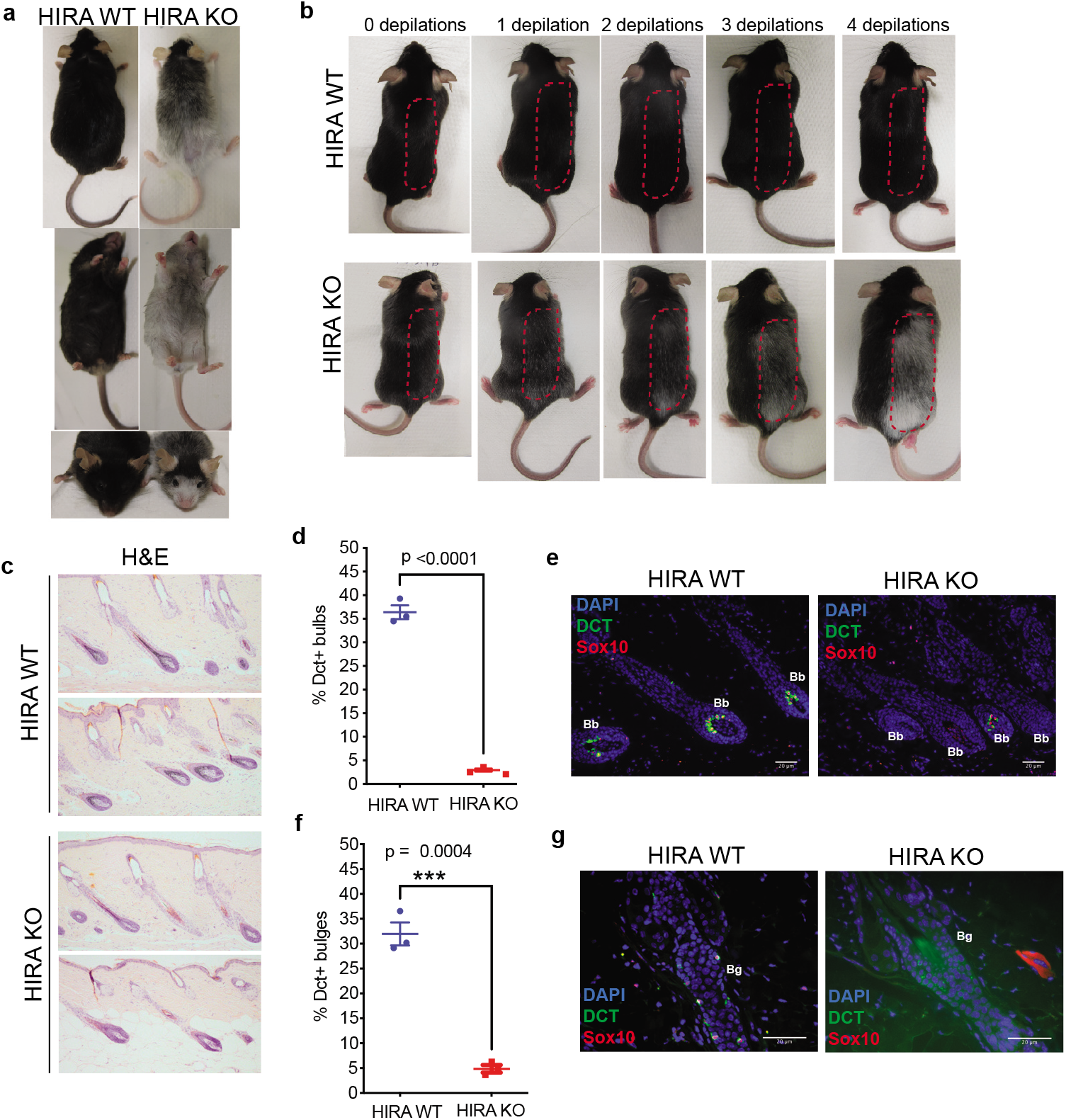
HIRA is required during embryonic development for McSC maintenance in adults and suppression of hair graying during aging. **a,** Back, belly and frontal images (top to bottom) of 20 month old mice showing premature hair graying in *TyrCre::Hira^fl/fl^* (HIRA KO) mice as compared to *TyrCre::Hira^wt/wt^* (HIRA WT) mice which retained their dark coat colour. **b,** Progressive hair graying and loss of pigment in the depilated side of the back skin in HIRA KO mice, while HIRA WT mice retained hair pigment. Images were taken before the first depilation and after full hair growth following each round of depilation, usually every 3 weeks. 4 rounds of depilation were performed in total over a 3 month period, starting with 8 week old mice; representative of n = 8 mice. **c,** H&E stain of longitudinal skin sections from 13 month old HIRA WT and HIRA KO mice. Skins were harvested 2 weeks following depilation to synchronise hair cycles. **d,** Quantification of differentiated melanocytes in the skins of mice in (**e**) by measuring the percentage of DCT/SOX10 positive hair bulbs. A total of 220-230 anagen hair bulbs were counted per genotype; n = 3 mice each. **e,** Representative images of hair follicles quantified in (**d**). Scale bar: 20 μm. **f**, Quantification of McSC content in the same mice in (**d**). A total of 300-400 hair follicles (anagen and telogen phases) were counted for presence of DCT/SOX10 positive cells in the bulge region. Scatter dot plot data in **d** and **f** were analysed using an unpaired *t-*test showing mean ± s.e.m. **g**, Representative images of bulges counted in (**f**). Scale bar: 20 μm.

To probe McSC function directly, we performed sequential rounds of depilation beginning on HIRA WT and KO mice at 8 weeks of age. Indicative of a McSC functional defect, the HIRA KO mice showed progressive but marked hair graying over 4 cycles of depilation (**Fig. 5b**). The effects of depilation were particularly marked in older mice. We aged HIRA WT and KO mice to 13 months and then again challenged the McSCs by depilation. Coat colour of these mice exhibited signs of a premature hair graying phenotype after only 1 cycle of depilation (**Supplementary Fig. 5d**), and H&E longitudinal skin sections showed loss of pigment in HIRA KO anagen hair follicles compared to HIRA WT (**Fig. 5c**). We quantified differentiated melanocytes and McSCs in these 13 month old mice, identified by DCT and SOX10 positive cells in the hair bulb and bulge, respectively. Strikingly, while in HIRA WT mice 36.4% ±2.5% of hair bulbs contained melanocytes, only 2.95% ±0.6% of hair bulbs contained melanocytes in HIRA KO mice (**Fig. 5d, e**). Moreover, the percentage McSC-containing bulges of telogen and anagen hair follicles of HIRA KO mice was 4.9% ±1.3%, significantly lower than that in in HIRA WT mice (32% ±4%) (**Fig. 5f, g**). Our results indicate that HIRA is required during embryonic development for McSC maintenance and suppression of hair greying during aging.

## Discussion

Previous human observational studies have shown that environmental conditions during embryonic development, such as a mother’s nutrition or exposure to toxins, can have longterm effects on health and aging of the offspring^75^. Consistent with these epidemiological observations, in this study we showed that a specific developmental abnormality occurring *in utero* results in barely perceptible defects at birth and young adulthood, but marked deficiencies under stress and/or during aging. Specifically, knockout of the gene encoding histone chaperone HIRA in migratory neural crest cells had a strong impact on melanoblast numbers early in mouse embryonic development, although the melanocytic cell number recovered by birth and function of melanocyte stem cells appeared normal in young adults. However, these cells failed to respond to growth signals under stress, and later in adult life McSCs were prematurely depleted and hair pigment was prematurely lost.

Our data in the melanocytic lineage support the essential role of HIRA for normal cell differentiation and development. Previous studies have shown that HIRA is required for gastrulation, patterning of meso-endodermal cells and embryonic viability^21,76^, and proper differentiation of ES cells^27^, muscle cells^77^, hematopoietic cells^28^, neurogenesis^31^ and oogenesis^25^. Extending these observations, we show here that HIRA is required for normal embryonic development of SOX10+ melanoblasts and expression of melanocytic genes, DCT and MITF. The decrease in SOX10+ melanoblasts was accompanied by a reciprocal increase in mesenchymal cells, including an abnormal population with features of EMT, perhaps indicative of a switch from a melanoblast to a mesenchymal phenotype. Consistent with this idea, other studies have previously shown a switch of melanocytic to mesenchymal lineages, particularly on downregulation of SOX10^78,79^, as is observed here. HIRA has been shown to physically interact with PAX3, a transcription factor known to play an essential role in melanoblast specification^80^. Whether the role of HIRA in the melanocytic lineage is dependent on H3.3 deposition or interaction with other proteins remains to be investigated.

Despite the consistent decrease in number of melanoblasts in E11.5-E15.5 HIRA KO embryos, newborn and young HIRA KO mice exhibited essentially normal numbers of melanoblasts, McSc and melanocytes, normal pigmentation and normally functioning McSC. Consistent with this, the melanoblast deficit in embryos showed a trend to a decrease from E11.5 to E15.5, although this was not significant at least with these cohort sizes. There are several possible mechanisms by which melanoblasts might recover their numbers during late embryogenesis. First, HIRA KO melanoblasts could compensate for their reduced number during early embryogenesis by increased proliferation in late embryogenesis. However, at E13.5 and E15.5 there was no difference in BrdU incorporation in WT and HIRA KO melanoblasts, suggesting that this is likely not the case, although we cannot exclude a late proliferative burst. Second, HIRA KO melanoblasts might compensate for their reduced number during early embryogenesis by decreased apoptosis in late embryogenesis. Consistent with this possibility, during normal embryogenesis a proportion of melanoblasts is culled by developmental apoptosis, ~0.5-1% melanoblasts at E12.5^81,82^. Third, HIRA KO mesenchymal-like cells might undergo a compensatory reversion back to melanoblasts, perhaps because the initial switch to the melanoblast program is incomplete. Consistent with this third possibility, the HIRA KO mesenchymal population contains a population that is distinct from HIRA WT and expresses genes indicative of EMT, perhaps suggesting only an immature “meta-stable” mesenchymal phenotype^83^.

Despite superficially normal numbers, phenotype and function of melanocytic cells in new born and young adult mice, several lines of evidence show a functional deficit in HIRA KO melanocytic cells. First, in *ex vivo* assays, HIRA KO cells from new born mice are unable to proliferate and differentiate into pigmented melanocytes. Second, KO of *Hira* suppresses the hyperpigmentation of *TyrNRasQ61K* mice, which we and others have previously shown is due to a marked post-natal proliferative expansion to generate an abnormal abundance of pigmented melanocytes in the dermis^72,73^. Third, adult HIRA KO mice turn prematurely grey after repeated rounds of depilation or physiological aging, accompanied by depletion of McSCs. Together, these results suggest that HIRA KO melanocytic cells, melanoblasts and McSCs, are impaired in their ability to sustainably proliferate and/or self renew when challenged. Importantly, this phenotype depended on embryonic inactivation of *Hira*, suggesting that the defect reflects aberrant programming during early stages of lineage development and/or is accumulated during embryogenesis. There are at least two likely explanations of this. First, DCT+ cells from HIRA KO P1 neonates contained significantly more TAFs than those from WT mice. This may result from increased proliferation, and hence telomere erosion and/or uncapping, during late embryogenesis as the HIRA KO melanoblasts recoup their numbers. Regardless of their origin, TAFs are often associated with cell senescence and aging^74,84–90^, including in skin melanocytes^74^. Therefore, the TAFs in neonate and young adult melanocytic cells might limit their proliferative/self-renewal capacity, ultimately leading to McSC exhaustion and premature hair greying. Second, histone H3.3, and by extension HIRA, is implicated in epigenetic memory through cell division^18–20^. Therefore, defects in HIRA-mediated histone H3.3 deposition during embryo development may translate into “silent” epigenetic defects in newborn mice, but progressive defects in epigenetic memory, gene expression and cell fate determination through sequential rounds of cell division during adulthood, and hence premature loss of McSCs and hair greying. Consistent with this model, hair bulb melanocytes are particularly enriched in histone H3.3 suggesting a specialized function for histone H3.3 in melanocytes.

Dysregulation of the SOX10-MITF axis contributes to melanoma. *MITF* and *SOX10* are both amplified and/or overexpressed in melanoma and are drivers of this disease^91^. Remarkably, *HIRA* is also altered in ~23% (mostly amplification/increased expression, no deletions (TCGA, cBioportal)) and overexpressed in ~15% of melanoma (TCGA, cBioportal). *HIRA* and mRNA encoding HIRA’s binding partner CABIN1 are co-overexpressed with *SOX10* in a significant proportion of melanoma. This suggests a potential oncogenic function for elevated HIRA expression. Based on findings reported here, upregulation of HIRA might confer a selective advantage during melanogenesis by upregulation of the SOX10-MITF signaling pathway.

In sum, these studies reveal a role for HIRA in proper differentiation and development of mouse embryo melanoblasts. This role for HIRA in promoting expression of melanoma oncogenes, MITF and SOX10, might account for *HIRA’s* overexpression in melanoma. However, these results also demonstrate that significant abnormalities in embryonic development can be superficially normalized by birth, only to manifest in later life as defects in maintenance of adult tissue function. These results highlight the potential connections between proper *in utero* development of tissue stem and progenitor populations and phenotypes characteristic of late life healthy aging.

## Methods

### Mice

*TryCre::Hira^fl/fl^* mice were generated by crossing *Hira^fl/fl^*^34^ with *TyrCre^53^* mice. They were then crossed with either *DCT::LacZ^55^* or *LSL-tdTomato^54^* reporter mice for melanoblast or melanocyte tracking experiments. Mice were genotyped both by PCR analysis and Transnetyx (www.transnetyx.com). *TyrCreERT2* mice^92,93^ were induced by applying 2 mg/ml Tamoxifen (Sigma) on shaved back skin of 6 week old mice once a day for 5 consecutive days. Depilations of back skin hair of adult mice was performed using Veet Easy-Gelwax strips under short anaesthesia with isofluorane. All experiments were carried out under the UK Home Office guidelines at the CRUK Beatson Institute biological services and research units.

### Fluorescence Immunohistochemistry

Shaved back skin from postnatal mice was fixed with 10% neutral buffered formalin (NBF), embedded in paraffin then cut longitudinally along the direction of hair growth. Embryos were harvested after a 2 hour BrdU (Sigma GERPN201) pulse to the mother then fixed in 10% NBF overnight (ON) and transversally cut and embedded back to back in paraffin with heads or tails down. For immunohistochemical (IHC) staining, paraffin sections were dewaxed in xylene (2 x 5 minutes) then rehydrated in serial ethanol washes (2 x 100%, 95% and 70%), 2 minutes each, followed by 5 minutes in water. Antigen retrieval was performed by boiling the samples in target retrieval solution (Dako S1699), pH = 6, for 25 minutes, followed by slow cooling at room temperature (RT). After rinsing in water, blocking solution (1% bovine serum albumin (BSA, ThermoFisher Scientific BPE1605-100) and 5 % donkey serum (Sigma D9663) in PBS) was added to tissue sections for 1 hour at RT, followed by ON incubation in a closed humid box at 4°C with primary antibodies (**Supplementary Table 9**) diluted in blocking solution containing 0.5% Tween-20 (Sigma P7949). The following day, tissue sections were washed 2 x 5 minutes with 0.5% Tween-20/PBS (TPBS), then incubated with secondary antibodies (**Supplementary Table 10**) diluted 1/500 in blocking solution with 0.5% Tween-20 for 40 minutes in the dark at RT. They were then washed 2 x 5 minutes with TPBS, stained with 1 μg/ml DAPI (Sigma 9542) and mounted in ProLong gold antifade (ThermoFisher Scientific P36934). Most images were taken using Nikon Eclipse 80i microscope fitted with a Hamamatsu ORCA-ER digital camera (C4742-80) or Nikon A1R confocal microscope.

### Immuno-FISH (TAF assay)

For FFPE tissues, immunohistochemistry was carried out following an overnight incubation with rabbit monoclonal anti-γH2AX (1:200, 9718; Cell Signalling), sections were incubated with a goat anti-rabbit biotinylated secondary antibody (1:200, PK-6101; Vector Labs) for 30 min at room temperature. Following three PBS washes, tissues were incubated with avidin DCS (1:500, SA-**A-2011-1**; Vector Labs) for another 30 min at room temperature. Sections were then washed three times in PBS blocked for 1h and incubated overnight with DCT (goat, Santa Cruz sc-10451, 1:200), sections were washed 3 times with PBS then incubated for 1h with a donkey anti-goat 647 (1:1000, **Catalog #** A-21447, invitrogen) followed by 3 PBS wash and cross-linked by incubation in 4% paraformaldehyde in PBS for 20 min. Sections were washed in PBS three times and then dehydrated in graded cold ethanol solutions (70, 90, 100%) for 3 min each. Tissues were then allowed to air-dry prior to being denatured in 10 μl of PNA hybridisation mix (70% deionized formamide (Sigma), 25 mM MgCl_2_, 1 M Tris pH 7.2, 5% blocking reagent (Roche) containing 2.5 μg/ml Cy-3-labelled telomere-specific (CCCTAA) peptide nucleic acid (Panagene), then denatured for 10 min at 80°C and then incubated for 2 h at room temperature in a dark humidified chamber to allow hybridization to occur. Sections were washed in 70% formamide in 2× SCC for 10 min, followed by a wash in 2× SSC for 10 min, and a PBS wash for 10 min. Tissues were then mounted using ProLong Gold Antifade Mountant with DAPI (Invitrogen). Sections were imaged using in-depth Z stacking (a minimum of 40 optical slices with 63 × objective) followed by Huygens (SVI) deconvolution.

### X-gal staining of embryos

Embryos carrying *DCT::LacZ* reporter gene were harvested at E11.5, E13.5 or E15.5, fixed in ice-cold 0.25% gluteraldehyde for 30 minutes and stained as previously described^71^. In brief, embryos were washed in ice-cold PBS then permeablised in a solution containing 0.02% NP-40, 2 mM MgCl_2_ and 0.01% sodium deoxycholate in PBS for 30 minutes at RT. Embryos were then stained with X-gal solution containing 1 mg/ml X-gal (Promega V3941), 0.01% sodium deoxycholate, 0.02% NP-40, 5 mM K_4_Fe(CN)_6_, 5 mM K_3_Fe(CN)_6_ for 3 nights (E15.5 and E13.5 embryos) or overnight (E11.5 embryos) at 4°C rotating in the dark. Stained embryos were then washed in PBS, post-fixed in 10% NBF ON at 4°C, then stored in 70% ethanol. Images were taken using Zeiss Stemi 2000-C dissection microscope fitted with Canon EOS Rebel 1000D camera, and stained cells were quantified using ImageJ.

### tdTomato positive cell sorting and single cell RNA sequencing

Embryos carrying tdTomato reporter were harvested at E15.5 and assessed for tdTomato positivity under a fluorescent microscope. Skin pieces from ears and tail were used for genomic DNA extraction and PCR analysis to determine *Hira^fl^* recombination using two forward primers 5’AATGGTGCTTGCT TTTGTGG3’ and 5’TGAAGGTATGGAGGACGCTG3’ and one reverse primer 5’GCATTAC TTAATCCCCAGATGC3’. Embryos were washed in ice-cold HBSS. Heads, limbs and tails were cut off and trunk skin was peeled using a scalpel and forceps on the lid of a petri-dish on ice. Individual skins were rinsed in HBSS then cut and digested in 300 μl of 0.2 U/ml dispase (Thermofisher Scientific 17105-041) and 0.5 mg/ml DNaseI (Sigma DN25) in HBSS for 45 minutes at 37°C. Ice-cold 5mM EDTA/PBS was then added to skin suspensions up to a volume of 1 ml to stop the digestion. The mix was passed through 18G followed by 21G needles for full dissociation, then filtered through pre-wet 50 μm filter membranes (Sysmex 04-0042-2317) into FACS tubes on ice. Cell suspensions were centrifuged at 300g for 5 minutes at 4°C then resuspended in 500 μl sorting buffer (1% FBS, 5mM EDTA and 25 mM Hepes in PBS). For sorting, at least one tdTomato^+^ *Hira^wt/wt^* and one *Hira^fl/fl^* individual skins from the same litter were used. tdTomato-skins were used as negative control for gating. 1 μg/ml DAPI was used as dead cell marker prior to sorting. tdTomato^+^ live cells were sorted using a BD FACSAria sorter, and 12000 to 20000 cells were collected per skin piece, corresponding to 1-2% of the whole cell suspension. Approximately 7000 to10000 cells were used for single-cell RNA isolation, cDNA synthesis and library preparation according to Chromium protocol (10x Genomics Chromium Single Cell 3’ Reagent kits 120237, 120236 and 120267). Libraries per experiment were pooled and sequenced on Illumina Hi-Seq4000. **Supplementary Table 11** shows details of the 3 WT and 3 KO biological replicates from 3 different embryos per genotype used for scRNAseq analysis. WT1 and KO1 are littermates and were processed simultaneously and pooled in the same sequencing lane. WT2, WT3 and KO2 were littermates, and KO3 was from a separate litter but had a simultaneous plug mating as the other 3 samples. The last 4 samples were processed simultaneously and sequenced in the same lane.

### scRNAseq analysis method

Initial data processing was performed using Cellranger version 2.2.0 (10X genomics). A novel mm10 reference genome was created that included tdTomato and Cre sequences using protocols provided by 10X genomics, then counts matrices of scRNAseq reads from each individual sample were created using this custom reference genome and the Cellranger ‘count’ function. All six samples were then combined into one experiment using the Cellranger ‘aggr’ function with the setting -normalize=mapped. Next, a counts matrix was created using the R packages dplyr and Seurat^59^, and filtered such that genes expressed in at least 3 cells, and cells with at least 100 expressed genes, were retained. The final matrix contained data for 18090 genes across 18453 cells, and was annotated with the sample ID, genotype and sex. To create feature plots, Seurat was used as follows. Firstly, the counts matrix was filtered to exclude cells with more than 40000 UMIs and more than 10% mitochondrial genes. The data were then log normalised, variable genes were identified using a logVMR dispersion function, and the data were scaled. Dimension reduction was performed using the ‘RunPCA’ and ‘RunTSNE’ functions, and feature plots were created using the ‘FeaturePlot’ function. Clusters were mapped to cell types using gene expression profiles.

Differential expression analysis of WT *vs* KO cells in the total cell population and the indicated subsets was performed using The R packages Scater, Scran and MAST^94,95^. Briefly, the annotated, filtered counts matrix was converted to a SingleCellExperiment object, then filtered to remove outliers based on the number of median absolute deviations (nmads) from the median as follows: cells with total counts more than 3 nmads from the median, cells with total features more than 4 nmads from the median, and cells with mitochondrial content more than 4 nmands from the median were dropped. Genes with an average expression <0.1 across all cells were then dropped. The resulting matrix (5985 cells by 5053 genes) was then normalised using the ‘computeSumfactors’ function, and variable genes were identified using a gamma-distributed GLM. Expression from the 4485 highly variable genes were then correlated, and heirarchical clustering performed using Ward’s clustering criterion. Gene expression in the resulting clusters was used to identify the indicated cell populations, and the dataset was trimmed to contain only subsets of clusters, as appropriate. Differential expression analysis was performed using a zero-inflated regression model from the package ‘MAST’, and the most significantly changed genes were noted.

### Melanoblast quantification from tdTomato populations

tdTomato+ skin cell suspensions from E15.5 embryos were prepared and genotyped as described above. After digestion and initial filtering step, cells were pelleted then resuspended in 2 ml ice-cold staining solution (0.5% BSA/PBS) and counted using a haemocytometer. Cells were pelleted and resuspended in 200 μl staining solution, and 1 μl of Fc block (BD Biosciences 553142) was added per tube for 10 minutes on ice. In addition to tubes corresponding to individual skins from WT and KO, controls were prepared corresponding to tdTomato-cell suspension (unstained), BV711 FMO (fluorescence minus one), tdTomato FMO. BV711 conjugated CD117 (c-kit) antibody (BioLegend 105835) was added to cell suspensions corresponding to WT, KO and tdTomato FMO tubes at 1 μg per million cells. An additional control corresponding to isotype control (BD Biosciences 563045) was also used. Cells were incubated with antibodies in the dark for 30 minutes on ice. 1 ml of staining solution was added per tube and the cells were pelleted and resuspended in sorting solution for FACS analysis as described above. BV711 positive cells within the tdTomato population corresponded to melanoblasts. Flow cytometry was performed using BD FACSAria. FlowJo was used for gating and cell quantification (**Supplementary Fig. 1**).

### Melb-a culture

Melb-a line^65^ was cultured in RPMI medium (Gibco) containing 10% FBS, 2 mM L-glutamine, 25 U/ml penicillin, and 25 μg/ml streptomycin, 20 ng/ml stem cell factor (SCF) (Sigma S9915) and 40 pM fibroblast growth factor 2 (Fgf2) (Sigma SRP4038). Cells were grown at 37°C with 5% CO2 and passaged when reaching 80-90% confluency.

### Plasmids, transfections and viral infections

LRMIP-shHira1 and LRMIP-shHira2^34^ were used to knockdown *Hira* in melb-a cells, with pLRMIP-shluc used as control. Lentiviruses were produced by transfecting 293T cells with 20 μg of hairpin plasmids, 8 μg psPAX2 (Addgene) and 4 μg pLP/VSVG (Invitrogen) using lipofectamine 2000. Following six hours, transfection medium was replaced with fresh culture medium (DMEM containing 10%FBS and 2mM L-glutamine). Virus-containing supernatants were collected 24 hours and 48 hours following transfections and were used to infect melb-a cells supplemented with 8 μg/ml polybrene (Millipore TR-1003-G) ON at 37°C. Successfully infected cells were selected using 1 μg/ml puromycin (Invivogen ant-pr) in melb-a culture medium.

### Primary melanocyte isolation and culture

Primary mouse melanocytes were isolated and cultured as previously described^71^. Back skin from 1, 2 or 3 day old pups was peeled and rinsed in PBS, then cut and incubated in collagenase I + IV (5 mg/ml, ThermoFisher Scientific 17100017 & 17104019) for 40 minutes at 37°C. Tail tips of pups were sent to Transnetyx for genotyping. After washing in HBSS (ThermoFisher Scientific 14065-049), skin pieces were incubated with cell dissociation buffer (ThermoFisher Scientific 13151014) for 10 minutes at 37°C. Skin pieces were then further dissociated using 18G and 21G needles then washed in HBSS. Finally, cells were resuspended and cultured in F-12 (Gibco) growth medium containing 10% fetal bovine serum (FBS), 200 nM12-tetradecanoylphorbol-12-acetate (TPA) and 100 μg/ml primocin (Invivogen ant-pm-1) at 37°C with 5% CO_2_. To select against fibroblasts and keratincytes, cultures were treated for 3 days a week with 50 μg/ml G418 (Formedium). Pure melanocyte cultures were established 6-8 weeks following initial isolation.

### Immunofluorescence and EdU labelling of cells

Cells were seeded on 13 mm cover glass prior to immunofluorescence (IF). Cells were washed in PBS then fixed in 4% paraformaldehyde for 10 minutes at RT. They were washed in PBS then permeablised and blocked in 3% BSA and 1% serum of the secondary antibody species in PBST (0.1% Triton-X in PBS). Cells were then incubated ON at 4°C in primary antibodies: DCT (goat, Santa Cruz sc-10451, 1:200); c-Kit (goat, R&D systems AF1356, 1:400); BIII tubulin (Tuj1, mouse, Promega G7121, 1:1000). After 3 x 5 minutes washes in PBST cells were incubated in secondary antibodies (**Supplementary Table 10**) for 1 hour at RT in the dark. Finally, cells were washed 3 x 5 minutes in PBST, stained with 1 μg/ml DAPI and mounted on glass slides in ProLong gold antifade. Images were taken using Nikon A1R confocal microscope.

For cell proliferation analysis by EdU labelling, cells were pulsed with 10 μM EdU for 2 hours (melb-a cells) or 48 hours (primary melanocytes) then fixed for IF. Click-iT EdU reaction mix was added to the cells following secondary antibody incubation and prior to DAPI staining as described by manufacturer’s protocol (ThermoFisher Scientific C10337 or C10339). EdU positive nuclei were quantified using ImageJ.

### Western Blot

Whole cell lysates were prepared by boiling and sheering cells in 1x Laemmeli sample buffer, and protein concentrations were measured using Qubit protein assay kit (ThermoFisher Scientific Q33212). 5 μg to 15 μg of proteins were separated on 4-15% SDS-PAGE gels (BioRAD 456-1084) and transferred onto PVDF membranes. Spectra multicolor broad range protein ladder (ThermoFisher Scientific 26634) used as a protein size marker Western blotting was performed as previously described^34^. Primary and secondary antibodies are listed in Supplementary **Tables 12** and **13** respectively.

### Statistics

Analysis was performed using GraphPad Prism (Version 9.0.1) and presented as mean ± s.e.m. Differences between two groups were assessed for statistical significance by an unpaired *t*-test. p values ≤ 0.05 were considered significant.

### Data Availability

scRNAseq data were deposited in the Gene Expression Omnibus (GEO) and can be accessed, using reviewer token **gjmdcmimzdollol,** through the following link: https://www.ncbi.nlm.nih.gov/geo/query/acc.cgi?acc=GSE132545. All raw data used to generate the results and figures of this study are available upon request. This includes scripts used for scRNAseq analysis, original IHC and IF images, quantification of dermal pigmentation, original Western blot scans and individual cell counts for melanoblasts, melanocytes and McSCs (DCT-LacZ, BrdU, EdU, DCT immunofluorescence, TAFs and flow cytometry).

## Acknowledgements

Work in the lab of PDA was supported by P01 AG031862-13 and from additional funding from CRUK Glasgow Centre (A25142) and Core Services at the Cancer Research UK Beatson Institute (A17196). LMM was funded by core CRUK grant A15673 and A24452. Work in the lab of JPM was supported by U24 CA220341, U24 CA194107, and U24 CA248457. ATW is supported by NIH grant T15LM011271. JFP and AL are supported by R01 AG069048, the Ted Nash Long Life Foundation and UL1 TR0002377 from the National Center for Advancing Translational Sciences (NCATS). We thank members of the Biological Services Unit at CRUK Beatson Institute for help in animal maintenance, members of Histology services, especially Colin Nixon, for help in mouse tissue processing, and Tom Gilbey for help in cell sorting. We thank Catherine Winchester for proofreading and editing and we thank all members of the Adams, Machesky and Insall labs for critical discussions.

## Author contributions

FJ-H performed the vast majority of the experiments. KS provided experimental, computational and intellectual assistance and advice and optimised melanoblast sorting and scRNAseq with FJ-H. KS, KG, ATW, NR performed single cell RNA-seq analysis. AL performed TAF assays. KK assisted with single cell RNA-seq. CR, JP, TSR, NF provided experimental support. KB advised on mouse experiments. JPM advised on single cell RNA-seq analysis. JFP advised on TAF assays. LMM assisted in direction of the study. PDA and FJ-H conceived and directed the study. FJ-H and PDA wrote the manuscript. All authors edited the manuscript.

---

**Supplementary Fig.1 – Gating strategy for tdTomato+ and c-KIT+ cell quantification.** Gating was performed with FlowJo software and used to quantify BV711-CD117 (c-KIT) positive melanoblasts from tdTomato positive populations of dissociated E15.5 trunk skin cell suspensions. FMO: fluorescence minus one.

**Supplementary Fig. 2 Hira is required for melanoblast development. a,** tdTomato fluorescence in FACS isolated cells (samples described in **Fig. 2a**). **b,** t-SNE plot generated using Seurat package showing 13 cell clusters (numbered 0-12) based on gene expression relationship from 3 HIRA WT and 3 HIRA KO tdTomato+ cells grouped together. **c,** Heat map displaying top 5 distinguishing genes of each of the 13 clusters in (**b**). **d-e,** t-SNE plots from combined 3 WT samples showing the expression of various melanoblast **(d)**, glial **(e)** and mesenchymal **(f)** genes in different clusters as viewed from Loupe Browser 4.2.0 by10xGenomics. Genes were taken from the lists in **Supplementary Tables 2-4**. **g,** 100% stacked column graph displaying the percentage of each cell type within the tdTomato population within each of the 6 samples described in **Fig. 2e**. **h,** Scatter dot plots showing the mean percentages of the major cell populations for each of 3 HIRA WT and 3 HIRA KO replicates. Data were analysed using an unpaired *t-*test showing mean ± s.e.m. **i,** GSEA showing KO cells enriched for the EMT signature relative to WT cells (NES=1.43, FDR q = 0.0081). **j-m** Western blots for various proteins using cell lysates harvested 10 days following lentiviral infections of melb-a cells with shLuc, shHira1 and shHira2. Multiple lanes for each represent independent replicates. GAPDH, Actin and LaminA/C were used as sample integrity controls.

**Supplementary Fig. 3. Knockout of *Hira* does not affect the distribution and number of melanocytes in young adult mice. a.** Mild difference in hair pigmentation in HIRA KO mice as compared to HIRA WT litter mates at P33 during the first real anagen. **b**. DCT/DAPI immunofluorescence and its corresponding phase contrast images showing the presence of pigmented melanocytes in growing hair bulbs of HIRA WT and HIRA KO mouse skin at P33. Scale bar: 20 μm.

**Supplementary Fig. 4. No effect after inducible knockout of *Hira* in differentiated melanocytes *ex-vivo* and *in vivo,* but suppression of hyperpigmented *TyrNras* phenotype by constitutive *Hira* knockout. a,** *Ex vivo* melanocyte cultures from *TyrCreERT2* system with *Hira^wt/wt^* or *Hira^fl/fl^*, obtained from P1 pups and induced with 4-hydroxytamoxifen (4-OHT) at the same time of seeding. Images were taken 3 weeks following culture and prior to any passage. **b,** UV image of a gel electrophoresis of the EtBr-stained products of a PCR testing recombination in induced *TyrCreERT2::Hira^fl/fl^* cells. 4-OHT treated *TyrCreERT2::Hira^wt/wt^* cells and *Hira^fl/fl^* cells (without *TyrCreERT2*) were used as floxed allele and recombination controls. Band sizes: 717 bp bp for WT allele, 910 bp for floxed allele and 467 bp for recombined allele. First lane: 10 kb ladder (O’GeneRuler Fermentas SM1173). Second lane: no template control (NTC). **c,** Western blot with various melanocytic markers in cells from (**a**) and biological replicates (WT1 & KO1 were littermates and WT2 & KO2 were littermates from a separate litter). **d,** DCT immunofluorescence (green) of induced *ex vivo TyrCreERT2::Hira^wt/wt^ and TyrCreERT2::Hira^fl/fl^* cells fixed after 2 weeks in culture. Scale bar: 25 μm. **e,** Images of a HIRA WT C57Bl6 mouse displaying coat and skin colour. **f,** Distribution of melanocytes (DCT positive cells in green) in the skins of NRAS-HIRA WT and NRAS-HIRA KO p11 pups. Scale bar: 30 μm.

**Supplementary Fig. 5. Inducible post-natal knockout of *Hira* does not lead to hair graying. a**, Representative images of *TyrCreERT2::Hira^wt/wt^ and TyrCreERT2::Hira^fl/fl^* mice 18 months following tamoxifen induction at 6 weeks of age. 3 mice per genotype were observed. **b,** Colocalisation of tdTomato (red) and DCT (green) showing melanocyte specific recombination in *TyrCreERT2* system both in the bulb (Bb) and McSCs (magenta arrow heads) of the bulge (Bg). Images representative of n=4 mice. Scale bar: 25 μm. **c,** More images showing recombination in *TyrCreERT2* system in bulge McSCs of telogen (top panel) and anagen (lower panel) hair follicles. Scale bars: 25 μM top; 50 μm bottom. **d,** 13 month old HIRA WT and HIRA KO mice after 1 cycle of depilation.

