## Supplementary data for "Inactivation of histone chaperone HIRA unmasks a link between normal embryonic development of melanoblasts and maintenance of adult melanocyte stem cells"

### Supplementary Fig. 1

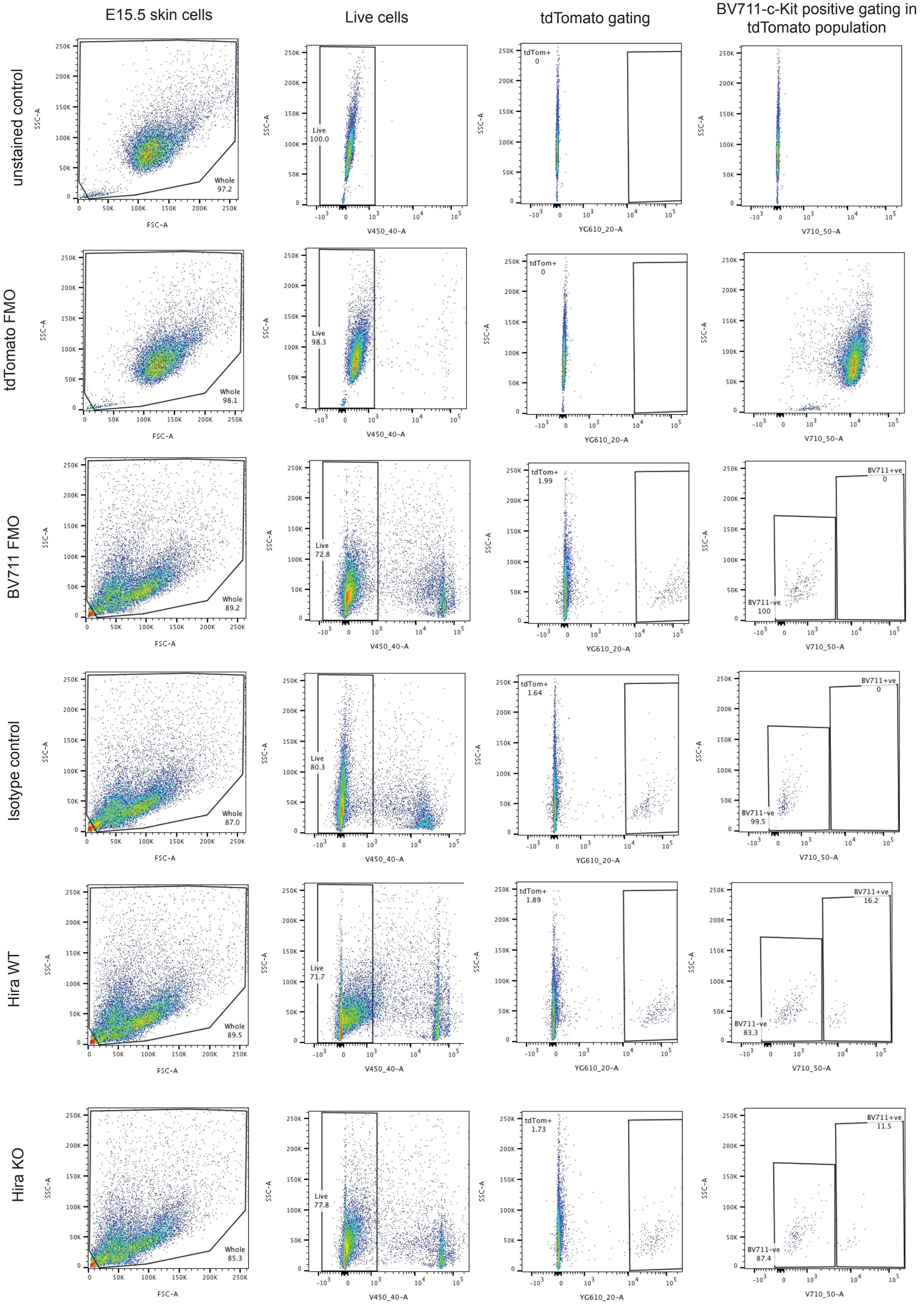

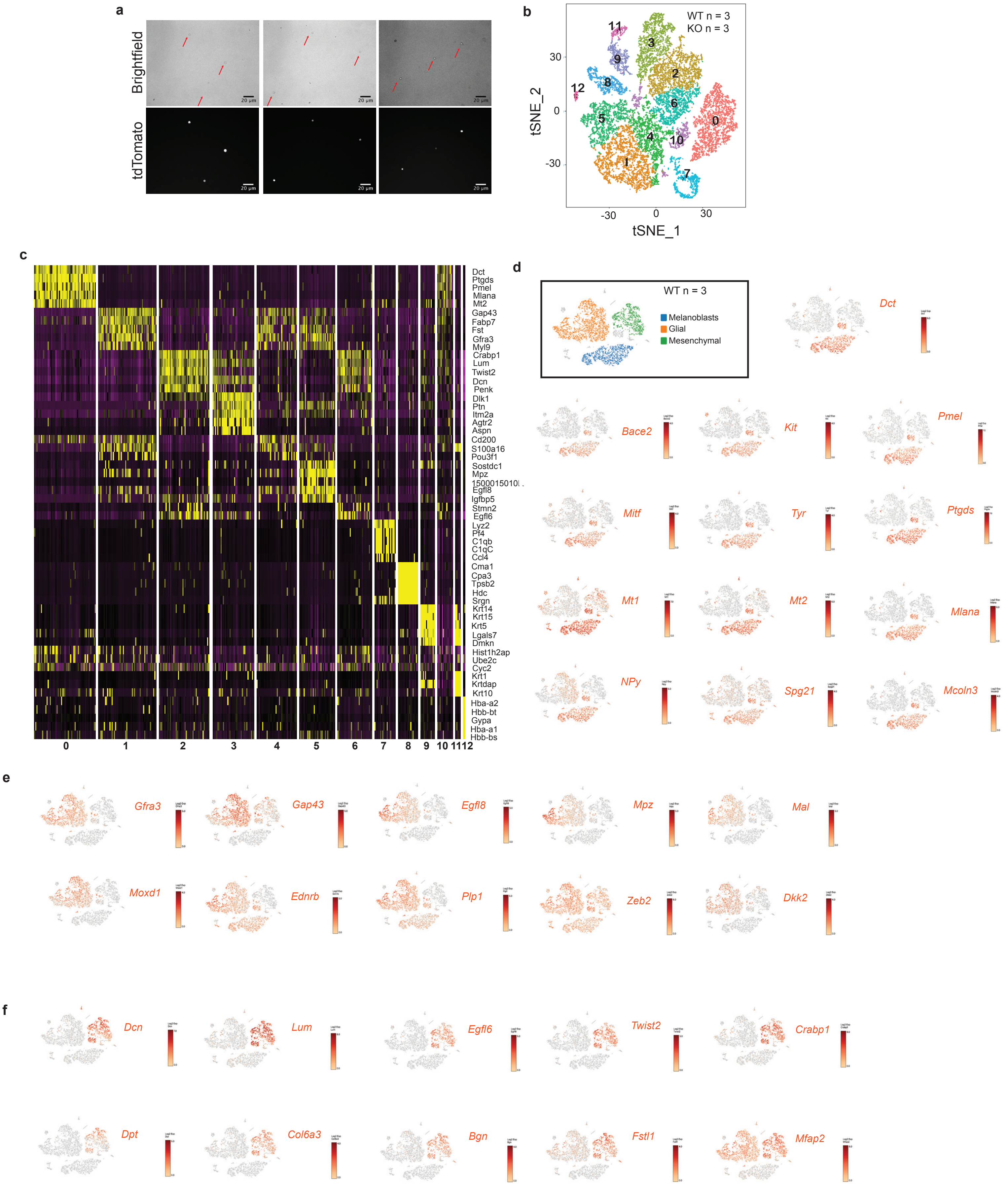
a
Brightfield
tdTomato
20 μm
b
WT n = 3
KO n = 3
tSNE_2
tSNE_1
c
Dct
Ptgds
Pmel
Mlana
Mt2
Gap43
Fabp7
Fst
Gfra3
Myl9
Crabp1
Lum
Twist2
Dcn
Penk
Dlk1
Ptn
Itm2a
Agtr2
Aspn
Cd200
S100a16
Pou3f1
Sostdc1
Mpz
Egfl8
Igfbp5
Stmn2
Egfl6
Lyz2
Pf4
C1qb
C1qC
Ccl4
Cma1
Cpa3
Tpsb2
Hdc
Srgn
Krt14
Krt15
Krt5
Lgals7
Dmkn
Hist1h2ap
Ube2c
Cyc2
Krt1
Krtdap
Krt10
Hba-a2
Hbb-bt
Gypa
Hba-a1
Hbb-bs
d
WT n = 3
Melanoblasts
Glial
Mesenchymal
Dct
Bace2
Kit
Pmel
Mitf
Tyr
Ptgds
Mt1
Mt2
Mlana
NPy
Spg21
Mcoln3
e
Gfra3
Gap43
Egfl8
Mpz
Mal
Moxd1
Ednrb
Plp1
Zeb2
Dkk2
f
Dcn
Lum
Egfl6
Twist2
Crabp1
Dpt
Col6a3
Bgn
Fstl1
Mfap2

Supplementary Fig. 2

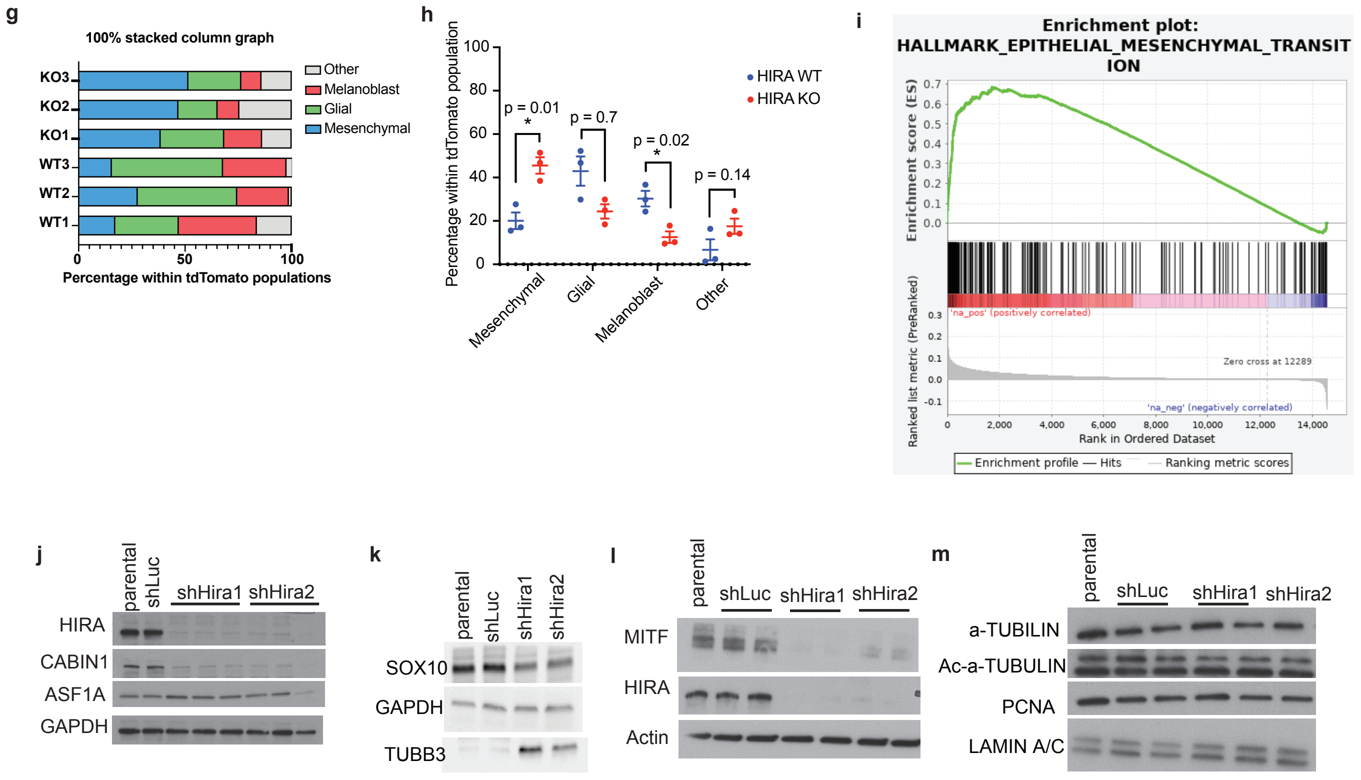

### Supplementary Fig. 3

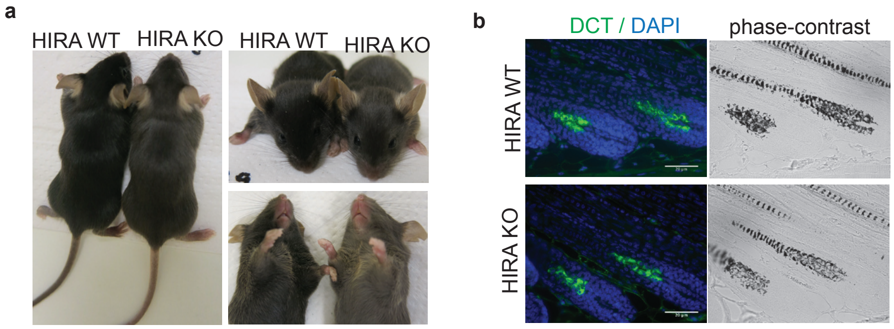

### Supplementary Fig. 4

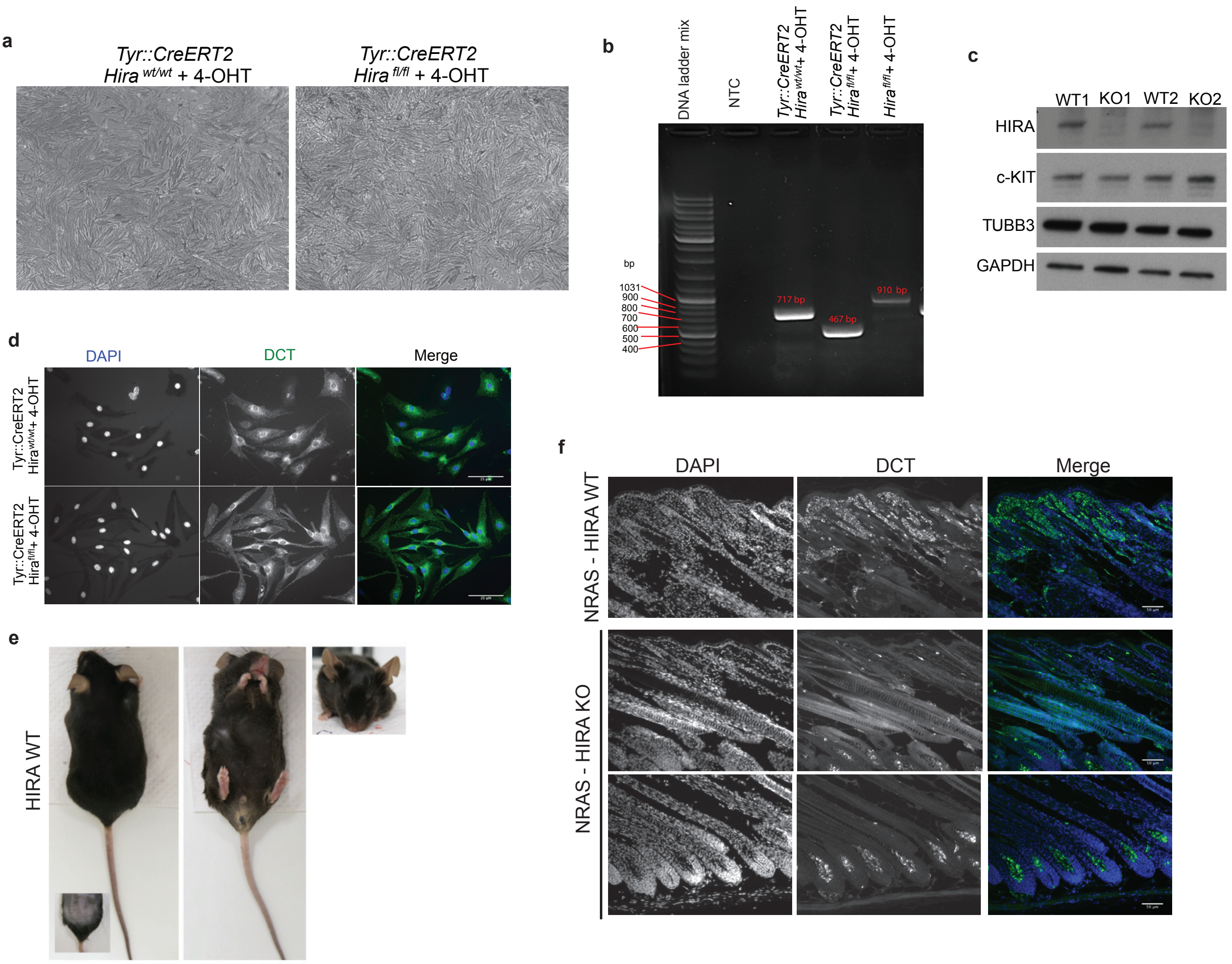

#### Supplementary Fig. 5

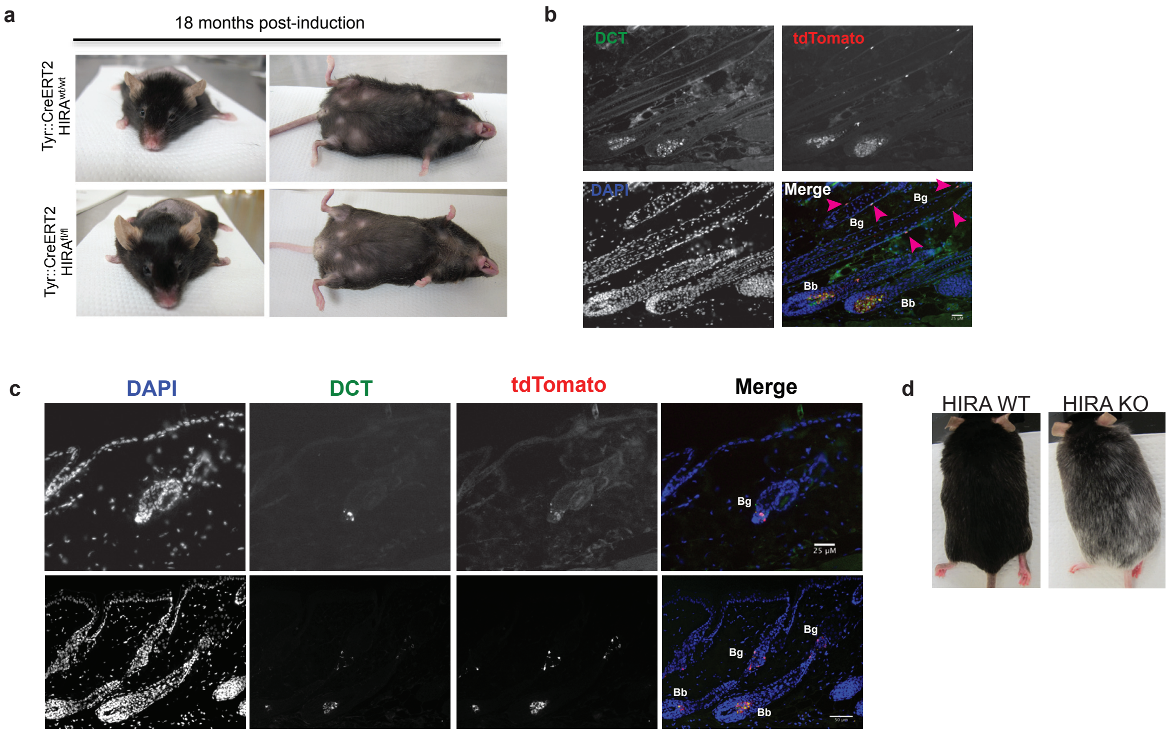

**Supplementary Table 1. Detailed information on the samples used for single cell RNA sequencing and the cells used for sequencing analysis**

| | WT1 | KO1 | WT2 | WT3 | KO2 | KO3 |
|---|---|---|---|---|---|---|
| Sex | F | M | M | F | F | F |
| Experiment | Chromium 1 | Chromium 1 | Chromium 2 | Chromium 2 | Chromium 2 | Chromium 2 |
| ***Cells*** | | | | | | |
| Estimated Number of Cells | 1,549 | 838 | 3,639 | 3,523 | 4,310 | 4,674 |
| Mean Reads per Cell | 108,584 | 229,719 | 26,246 | 20,047 | 21,854 | 18,957 |
| Median Genes per Cell | 1,483 | 1,791 | 1,202 | 1,536 | 1,732 | 1,772 |
| Total Genes Detected | 16,571 | 16,706 | 16,703 | 17,023 | 18,370 | 18,528 |
| Median UMI Counts per Cell | 3,955 | 5,195 | 2,686 | 3,955 | 4,958 | 4,908 |
| ***Sequencing*** | | | | | | |
| Number of Reads | 168,198,042 | 192,504,564 | 95,512,423 | 70,628,024 | 94,194,137 | 88,691,811 |
| Valid Barcodes | 98.40% | 98.40% | 98.50% | 98.20% | 98.40% | 98.40% |
| Reads Mapped Confidently to Transcriptome | 51.40% | 56.60% | 62.50% | 63.00% | 61.90% | 59.70% |
| Reads Mapped Confidently to Exonic Regions | 53.60% | 58.80% | 66.20% | 66.50% | 65.50% | 63.10% |
| Reads Mapped Confidently to Intronic Regions | 10.50% | 11.30% | 12.70% | 13.60% | 11.20% | 12.50% |
| Reads Mapped Confidently to Intergenic Regions | 3.00% | 2.80% | 3.60% | 3.20% | 3.70% | 3.50% |
| Reads Mapped Antisense to Gene | 4.30% | 3.80% | 1.90% | 1.60% | 1.70% | 1.60% |
| Sequencing Saturation | 90.90% | 94.70% | 76.70% | 57.50% | 52.70% | 45.70% |
| Q30 Bases in Barcode | 98.00% | 98.00% | 98.10% | 98.00% | 98.10% | 98.00% |
| Q30 Bases in RNA Read | 83.90% | 83.20% | 78.20% | 76.60% | 76.40% | 74.30% |
| Q30 Bases in UMI | 98.30% | 98.30% | 97.90% | 97.80% | 97.80% | 97.80% |

**Supplementary Table 2. Top melanoblast distinguishing genes within tdTomato population from 3 WT embryos generated with MAST against all tdTomato population.**

| Rank | Gene name | Log fold change | p-value |
|---|---|---|---|
| 1 | Dct | 2.74 | 0 |
| 2 | Ptgds | 2.56 | 0 |
| 3 | Mt1 | 2.50 | 0 |
| 4 | Pmel | 1.93 | 0 |
| 5 | Mt2 | 1.69 | 0 |
| 6 | Mlana | 1.31 | 0 |
| 7 | Phlda1 | 1.26 | 2.78E-290 |
| 8 | Sat1 | 1.23 | 4.46E-256 |
| 9 | Cyb5a | 1.20 | 0 |
| 10 | Lmo4 | 0.89 | 1.16E-179 |
| 11 | Syngr1 | 0.87 | 0 |
| 12 | Npy | 0.84 | 4.25E-200 |
| 13 | Sox10 | 0.83 | 0 |
| 14 | Apoe | 0.81 | 6.05E-169 |
| 15 | Sdcbp | 0.79 | 8.34E-188 |
| 16 | Vim | 0.76 | 1.07E-135 |
| 17 | Deb1 | 0.76 | 1.63E-185 |
| 18 | Mif | 0.73 | 3.95E-139 |
| 19 | Cd63 | 0.73 | 8.63E-82 |
| 20 | Slc24a5 | 0.70 | 1.34E-154 |
| 21 | Mcoln3 | 0.70 | 2.57E-306 |
| 22 | Gstp1 | 0.69 | 4.76E-140 |
| 23 | Fabp5 | 0.67 | 4.11E-91 |
| 24 | Atp6v1e1 | 0.65 | 2.42E-164 |
| 25 | 2700094K13Rik | 0.64 | 1.91E-102 |
| 26 | Mitf | 0.64 | 3.06E-255 |
| 27 | Prdx1 | 0.63 | 7.84E-159 |
| 28 | Tyr | 0.60 | 4.27E-256 |
| 29 | Npm1 | 0.58 | 6.44E-124 |
| 30 | H2afz | 0.58 | 5.07E-54 |
| 31 | Kit | 0.56 | 1.73E-238 |
| 32 | Bri3 | 0.55 | 1.57E-110 |
| 33 | Pax3 | 0.55 | 6.35E-161 |
| 34 | Slc25a5 | 0.54 | 1.25E-85 |
| 35 | B2m | 0.54 | 1.51E-88 |
| 36 | Atp6v1g1 | 0.54 | 1.70E-89 |
| 37 | Cotl1 | 0.53 | 6.61E-111 |
| 38 | Syt4 | 0.51 | 3.50E-204 |
| 39 | Idh2 | 0.49 | 3.78E-84 |
| 40 | Tm4sf1 | 0.49 | 2.74E-147 |
| 41 | Cfl1 | 0.48 | 2.33E-90 |
| 42 | Npc2 | 0.47 | 2.33E-76 |
| 43 | Spg21 | 0.47 | 6.64E-138 |
| 44 | Actg1 | 0.47 | 7.07E-70 |
| 45 | Xist | 0.46 | 3.00E-38 |
| 46 | H3f3a | 0.46 | 1.48E-90 |
| 47 | Cdk2 | 0.45 | 5.59E-127 |
| 48 | St3gal6 | 0.45 | 3.96E-194 |
| 49 | Bace2 | 0.44 | 1.99E-189 |
| 50 | Rpl22l1 | 0.42 | 1.20E-62 |

**Supplementary Table 3. Top glial distinguishing genes within tdTomato population from 3 WT embryos generated with MAST against all tdTomato population.**

| Rank | Gene name | Log fold change | p-value |
|---|---|---|---|
| 1 | Gfra3 | 1.38 | 0 |
| 2 | Dbi | 1.24 | 5.84E-265 |
| 3 | Gap43 | 1.20 | 1.50013152046778e-319 |
| 4 | Sparc | 1.18 | 1.10E-181 |
| 5 | Fabp7 | 1.17 | 1.83E-298 |
| 6 | Fst | 1.05 | 0 |
| 7 | Sox10 | 1.05 | 0 |
| 8 | Anxa2 | 0.79 | 8.55E-154 |
| 9 | Postn | 0.79 | 4.01E-235 |
| 10 | Plp1 | 0.77 | 5.92E-216 |
| 11 | Arpc1b | 0.75 | 1.95E-129 |
| 12 | Marcks | 0.74 | 1.75E-152 |
| 13 | Zeb2 | 0.71 | 1.77E-150 |
| 14 | Cryab | 0.71 | 3.07E-244 |
| 15 | Prss23 | 0.70 | 5.75E-241 |
| 16 | Cxcl12 | 0.70 | 1.77E-221 |
| 17 | Rbp1 | 0.70 | 5.11E-177 |
| 18 | Tuba1a | 0.69 | 1.25E-120 |
| 19 | Egfl8 | 0.67 | 5.36E-242 |
| 20 | Myl9 | 0.66 | 5.48E-244 |
| 21 | Tagln2 | 0.65 | 1.28E-128 |
| 22 | Cnn3 | 0.63 | 1.24E-125 |
| 23 | S100a16 | 0.59 | 2.89E-190 |
| 24 | Tmsb4x | 0.59 | 4.94E-69 |
| 25 | Anxa5 | 0.58 | 1.74E-112 |
| 26 | Cald1 | 0.56 | 1.95E-120 |
| 27 | Gpm6b | 0.56 | 9.89E-163 |
| 28 | Moxd1 | 0.55 | 4.48E-232 |
| 29 | Mef2c | 0.54 | 9.33E-96 |
| 30 | Sema3c | 0.52 | 7.78E-205 |
| 31 | Sep-15 | 0.52 | 2.63E-104 |
| 32 | Cox7c | 0.51 | 3.92E-129 |
| 33 | Mpz | 0.51 | 1.71E-165 |
| 34 | Timp3 | 0.50 | 2.73E-156 |
| 35 | Col4a1 | 0.50 | 9.44E-150 |
| 36 | Mest | 0.50 | 9.16E-126 |
| 37 | Cadm1 | 0.50 | 2.07E-132 |
| 38 | Mal | 0.49 | 7.78E-205 |
| 39 | Ednrb | 0.49 | 6.74E-95 |
| 40 | Kctd12 | 0.49 | 1.07E-123 |
| 41 | Dkk2 | 0.49 | 4.65E-170 |
| 42 | Mfap2 | 0.48 | 1.27E-141 |
| 43 | Fign | 0.48 | 1.36E-189 |
| 44 | Psat1 | 0.48 | 4.40E-113 |
| 45 | Tubb5 | 0.48 | 2.30E-64 |
| 46 | Foxp1 | 0.47 | 1.83E-117 |
| 47 | Gm29865 | 0.46 | 2.54E-149 |
| 48 | Malat1 | 0.46 | 1.59E-50 |
| 49 | Dst | 0.46 | 1.11E-149 |
| 50 | Pdgfa | 0.45 | 1.36E-161 |

**Supplementary Table 4. Top mesenchymal distinguishing genes within tdTomato population from 3 WT embryos generated using 10x Genomics Loupe browser.**

| Rank | Gene name | Log2 fold change | p-value |
|---|---|---|---|
| 1 | Dpt | 7.4016712 | 4.64E-121 |
| 2 | Egfl6 | 6.966488353 | 4.96E-139 |
| 3 | Dcn | 6.923427519 | 5.40E-163 |
| 4 | Lum | 6.842311174 | 5.40E-163 |
| 5 | Irx1 | 6.649462121 | 1.19E-109 |
| 6 | Twist2 | 6.449051683 | 1.05E-136 |
| 7 | C1qtnf2 | 6.404596077 | 7.52E-112 |
| 8 | Crabp1 | 6.371604798 | 7.47E-142 |
| 9 | Penk | 6.010439479 | 3.66E-117 |
| 10 | Dlk1 | 5.3126903 | 3.61E-101 |
| 11 | Col6a3 | 5.164827578 | 1.16E-92 |
| 12 | Bgn | 4.867429614 | 1.72E-84 |
| 13 | Prrx1 | 4.614590139 | 7.35E-78 |
| 14 | Col6a2 | 4.427982216 | 2.92E-78 |
| 15 | Kdelr3 | 3.850736578 | 7.88E-58 |
| 16 | Col6a1 | 3.784852504 | 4.94E-60 |
| 17 | Col1a1 | 3.395340797 | 2.85E-50 |
| 18 | Fstl1 | 3.174363647 | 2.81E-44 |
| 19 | Pcolce | 3.07675508 | 5.94E-41 |
| 20 | H19 | 2.998900806 | 1.60E-37 |
| 21 | P4ha2 | 2.838027938 | 2.12E-34 |
| 22 | Col3a1 | 2.816497294 | 6.91E-36 |
| 23 | Wif1 | 2.730979653 | 3.45E-30 |
| 24 | Col1a2 | 2.722140223 | 1.63E-33 |
| 25 | Cyr61 | 2.626555871 | 1.56E-29 |
| 26 | Rcn3 | 2.520898778 | 3.41E-29 |
| 27 | Tnfaip6 | 2.384903617 | 5.89E-24 |
| 28 | Meg3 | 2.273228891 | 2.05E-23 |
| 29 | Col5a2 | 2.206060721 | 2.59E-21 |
| 30 | Mfap2 | 1.460158406 | 3.45E-10 |
| 31 | Cd24a | 1.346838699 | 1.10E-07 |
| 32 | Nrn1 | 1.275269989 | 6.34E-07 |
| 33 | Hmgn3 | 1.224374348 | 3.82E-07 |
| 34 | Itih5 | 1.203483892 | 2.66E-06 |
| 35 | Ptn | 1.012014172 | 0.000155397 |

**Supplementary Table 5. Number and percentage of cells of the main clusters within the tdTomato population of 3 WT and 3 HIRA KO samples.**

| | Total number of cells per Sample | | | | | | Overall | |
|---|---|---|---|---|---|---|---|---|
| **Cellular Subset** | WT1 | WT2 | WT3 | KO1 | KO2 | KO3 | WT all | KO all |
| **All tdTomato** | 1534 | 3604 | 3517 | 814 | 4285 | 4644 | 8655 | 9743 |
| **Glial** | 458 | 1685 | 1837 | 243 | 789 | 1155 | 3980 | 2187 |
| **Mesenchymal** | 262 | 996 | 545 | 313 | 2004 | 2388 | 1803 | 4705 |
| **Melanoblast** | 563 | 875 | 1049 | 145 | 438 | 442 | 2487 | 1025 |
| **Other** | 251 | 48 | 86 | 113 | 1054 | 659 | 385 | 1826 |
| | **Percentage of cells in tdTomato population** | | | | | | | |
| **Glial** | 29.86% | 46.75% | 52.23% | 29.85% | 18.41% | 24.87% | 45.98% | 22.45% |
| **Mesenchymal** | 17.08% | 27.64% | 15.50% | 38.45% | 46.77% | 51.42% | 20.83% | 48.29% |
| **Melanoblast** | 36.70% | 24.28% | 29.83% | 17.81% | 10.22% | 9.52% | 28.73% | 10.52% |
| **Other** | 16.36% | 1.33% | 2.45% | 13.88% | 24.60% | 14.19% | 4.45% | 18.74% |

**Supplementary Table 6. Differentially expressed (DE) genes in HIRA KO *vs* HIRA WT melanoblasts calculated using MAST R package. Top 50 upregulated and top 50 downregulated genes are shown.**

| Melanoblast upregulated DE genes KO vs WT | | | | Melanoblast downregulated DE genes KO vs WT | | | |
|---|---|---|---|---|---|---|---|
| **Rank** | **Gene name** | **Log fold change** | **p-value** | **Rank** | **Gene name** | **Log fold change** | **p-value** |
| 1 | Fabp5 | 0.86 | 1.54566E-67 | 1 | Npy | -0.86 | 5.04E-52 |
| 2 | Xist | 0.61 | 1.14E-18 | 2 | Fabp7 | -0.54 | 2.50E-32 |
| 3 | Hist1h2ap | 0.51 | 7.74E-09 | 3 | Phlda1 | -0.41 | 1.12E-24 |
| 4 | AY036118 | 0.50 | 6.74E-28 | 4 | Cyb5a | -0.40 | 4.31E-27 |
| 5 | Pkm | 0.50 | 1.33E-25 | 5 | Vim | -0.39 | 6.74E-28 |
| 6 | Sparc | 0.50 | 4.17343E-22 | 6 | Sat1 | -0.36 | 1.34E-19 |
| 7 | Ctsl | 0.47 | 7.8799E-23 | 7 | Gm11808 | -0.34 | 1.46E-28 |
| 8 | Pgam1 | 0.46 | 1.09E-23 | 8 | Crip2 | -0.34 | 6.74E-28 |
| 9 | Gm42418 | 0.44 | 4.54E-20 | 9 | Lgals1 | -0.33 | 1.04E-10 |
| 10 | Tubb3 | 0.42 | 1.42E-34 | 10 | Olfm1 | -0.32 | 2.94E-25 |
| 11 | H2afz | 0.41 | 9.37E-17 | 11 | Rps27rt | -0.31 | 2.19E-28 |
| 12 | Ldha | 0.41 | 8.45E-16 | 12 | Ptgds | -0.30 | 2.47E-10 |
| 13 | Hsp90b1 | 0.41 | 5.73E-17 | 13 | Gm2000 | -0.29 | 7.45E-25 |
| 14 | H1f0 | 0.40 | 8.68E-20 | 14 | Tm4sf1 | -0.29 | 5.72E-15 |
| 15 | Marcksl1 | 0.40 | 3.34E-19 | 15 | Rpl36 | -0.28 | 6.86E-30 |
| 16 | Mif | 0.40 | 1.72E-23 | 16 | Smpdl3a | -0.28 | 4.41E-19 |
| 17 | Hmgb2 | 0.40 | 1.20E-09 | 17 | Rpl10 | -0.26 | 8.42E-25 |
| 18 | Col1a2 | 0.39 | 3.30E-11 | 18 | Emp3 | -0.26 | 1.64E-16 |
| 19 | Gnas | 0.38 | 3.64E-21 | 19 | Gm8730 | -0.26 | 5.25E-26 |
| 20 | Sub1 | 0.38 | 1.59E-27 | 20 | Lmo4 | -0.25 | 4.40E-15 |
| 21 | Col3a1 | 0.38 | 1.57E-14 | 21 | Dct | -0.25 | 4.74E-04 |
| 22 | Plgrkt | 0.37 | 1.72E-28 | 22 | Sox10 | -0.25 | 6.63E-18 |
| 23 | Col1a1 | 0.36 | 2.26E-15 | 23 | Rexo2 | -0.25 | 5.61E-25 |
| 24 | Pmel | 0.36 | 1.55E-07 | 24 | Cox20 | -0.24 | 1.67E-22 |
| 25 | Lum | 0.35 | 1.39E-14 | 25 | Chmp2a | -0.24 | 2.65E-22 |
| 26 | Eno1 | 0.34 | 6.78E-15 | 26 | Rpl41 | -0.23 | 1.49E-33 |
| 27 | Dcn | 0.34 | 3.03E-10 | 27 | Bmyc | -0.22 | 2.23E-22 |
| 28 | Nap1l1 | 0.34 | 2.85E-15 | 28 | Wdr89 | -0.22 | 2.88E-27 |
| 29 | Dlk1 | 0.32 | 2.98E-16 | 29 | Cyp2j6 | -0.22 | 1.89E-17 |
| 30 | Serpinh1 | 0.31 | 9.47E-12 | 30 | Slc26a7 | -0.22 | 1.59E-18 |
| 31 | Cbx3 | 0.31 | 1.42E-14 | 31 | Fau | -0.22 | 1.13E-18 |
| 32 | Psmd8 | 0.31 | 1.08E-15 | 32 | Gsta1 | -0.21 | 2.05E-15 |
| 33 | Rbm8a | 0.31 | 1.28E-15 | 33 | Rpl23a-ps3 | -0.21 | 1.16E-22 |
| 34 | Aldoa | 0.30 | 1.60E-10 | 34 | Gm9843 | -0.21 | 5.30E-21 |
| 35 | Snhg9 | 0.30 | 2.96E-23 | 35 | B2m | -0.21 | 1.18E-19 |
| 36 | Cadm1 | 0.30 | 9.12E-25 | 36 | Rplp1 | -0.20 | 1.74E-17 |
| 37 | Gapdh | 0.29 | 1.68E-11 | 37 | 1700086L19Rik | -0.20 | 1.01E-13 |
| 38 | Cma1 | 0.29 | 4.61E-43 | 38 | Sdpr | -0.20 | 3.59E-14 |
| 39 | Meg3 | 0.29 | 4.81E-08 | 39 | Nkain4 | -0.20 | 6.81E-17 |
| 40 | Calr | 0.29 | 2.99E-12 | 40 | Lima1 | -0.20 | 2.22E-20 |
| 41 | Hspd1 | 0.28 | 2.17E-12 | 41 | Ndufb10 | -0.19 | 4.91E-28 |
| 42 | Birc5 | 0.28 | 1.35E-12 | 42 | Plagl1 | -0.19 | 3.59E-12 |
| 43 | Crabp1 | 0.28 | 5.99E-10 | 43 | Actb | -0.19 | 5.40E-03 |
| 44 | Mdk | 0.28 | 1.16E-14 | 44 | Mgll | -0.19 | 4.60E-14 |
| 45 | Top2a | 0.28 | 7.43E-09 | 45 | Eci1 | -0.18 | 4.68E-19 |
| 46 | Minos1 | 0.28 | 3.02E-16 | 46 | Tagln2 | -0.18 | 6.62E-22 |
| 47 | Ran | 0.28 | 1.99E-11 | 47 | Lmna | -0.18 | 2.38E-16 |
| 48 | Arf1 | 0.28 | 1.70E-16 | 48 | Gm9493 | -0.17 | 1.70E-23 |
| 49 | Tma7 | 0.27 | 1.66E-12 | 49 | Hypk | -0.17 | 1.60E-18 |
| 50 | Hspa5 | 0.27 | 1.27E-11 | 50 | 2410015M20Rik | -0.17 | 4.65E-24 |

**Supplementary Table 7. Differentially expressed (DE) genes in HIRA KO *vs* HIRA WT glial cells calculated using MAST R package. Top 50 upregulated and top 50 downregulated genes are shown.**

| Glial upregulated DE genes KO vs WT | | | | Glial downregulated DE genes KO vs WT | | | |
|---|---|---|---|---|---|---|---|
| **Rank** | **Gene name** | **Log fold change** | **p-value** | **Rank** | **Gene name** | **Log fold change** | **p-value** |
| **1** | Lyve1 | NA | 5.18E-12 | **1** | Mfap2 | -0.59 | 6.93E-57 |
| **2** | Xist | 0.95 | 1.13E-50 | **2** | Fst | -0.55 | 3.93E-35 |
| **3** | Kctd12 | 0.58 | 2.47E-39 | **3** | Crip1 | -0.54 | 3.03E-25 |
| **4** | Tmsb4x | 0.49 | 1.13E-43 | **4** | Anxa2 | -0.50 | 4.84E-55 |
| **5** | H1f0 | 0.46 | 2.63E-28 | **5** | S100a6 | -0.37 | 4.78E-19 |
| **6** | Gm42418 | 0.46 | 3.59E-26 | **6** | Rps27rt | -0.37 | 6.48E-32 |
| **7** | AY036118 | 0.46 | 7.83E-30 | **7** | Crip2 | -0.35 | 6.52E-30 |
| **8** | Arpc1a | 0.41 | 3.03E-25 | **8** | Prss23 | -0.34 | 2.73E-19 |
| **9** | Fabp5 | 0.41 | 9.57E-22 | **9** | 1500015O10Rik | -0.34 | 1.05E-21 |
| **10** | Sub1 | 0.41 | 1.35E-30 | **10** | Id2 | -0.33 | 1.34E-16 |
| **11** | Pls3 | 0.38 | 4.54E-29 | **11** | Gfra3 | -0.32 | 4.89E-27 |
| **12** | Gnas | 0.34 | 2.98E-21 | **12** | Cox20 | -0.31 | 2.40E-28 |
| **13** | Marcksl1 | 0.34 | 1.13E-17 | **13** | 2410015M20Rik | -0.30 | 8.09E-26 |
| **14** | Tubb3 | 0.33 | 2.86E-38 | **14** | Rpl10 | -0.29 | 3.09E-29 |
| **15** | Hmgb2 | 0.33 | 3.14E-09 | **15** | Fabp7 | -0.28 | 3.26E-19 |
| **16** | Ccnd2 | 0.32 | 6.45E-14 | **16** | Prdx6 | -0.27 | 1.71E-20 |
| **17** | Mdk | 0.32 | 2.71E-15 | **17** | Rplp1 | -0.26 | 1.41E-27 |
| **18** | H2afz | 0.32 | 3.04E-09 | **18** | Ndufb10 | -0.26 | 7.66E-24 |
| **19** | Id3 | 0.31 | 2.40E-12 | **19** | Selm | -0.26 | 1.01E-20 |
| **20** | Akap12 | 0.31 | 3.52E-34 | **20** | Lims2 | -0.26 | 1.44E-20 |
| **21** | H3f3a | 0.29 | 5.52E-24 | **21** | Cryab | -0.25 | 7.05E-16 |
| **22** | Stmn1 | 0.28 | 3.36E-10 | **22** | Cd81 | -0.25 | 1.63E-18 |
| **23** | Dpysl3 | 0.28 | 2.61E-23 | **23** | Apoe | -0.25 | 5.23E-07 |
| **24** | Bsg | 0.27 | 1.03E-13 | **24** | Grb14 | -0.24 | 1.80E-12 |
| **25** | Actg1 | 0.27 | 7.22E-16 | **25** | Rpl36 | -0.24 | 3.81E-22 |
| **26** | Pabpc1 | 0.26 | 1.54E-12 | **26** | Rpl35 | -0.24 | 8.77E-28 |
| **27** | Cst3 | 0.26 | 6.58E-11 | **27** | Nxf1 | -0.23 | 6.65E-21 |
| **28** | Serbp1 | 0.26 | 3.13E-11 | **28** | Aqp1 | -0.23 | 3.83E-17 |
| **29** | Tmem176b | 0.26 | 1.06E-18 | **29** | Fau | -0.22 | 6.45E-19 |
| **30** | Hmga2 | 0.26 | 4.55E-23 | **30** | Lmna | -0.21 | 2.46E-16 |
| **31** | Lmo4 | 0.26 | 4.43E-10 | **31** | Hypk | -0.21 | 8.65E-19 |
| **32** | Hsp90aa1 | 0.25 | 4.08E-09 | **32** | Emp2 | -0.21 | 6.96E-15 |
| **33** | Dync1i2 | 0.25 | 1.03E-10 | **33** | Zcchc12 | -0.21 | 9.90E-22 |
| **34** | Cbx3 | 0.25 | 3.21E-08 | **34** | Gm11808 | -0.21 | 3.17E-19 |
| **35** | Hmgn1 | 0.24 | 2.11E-10 | **35** | Gm2000 | -0.21 | 6.96E-15 |
| **36** | Atp5k | 0.24 | 1.78E-11 | **36** | Atp5d | -0.20 | 6.10E-21 |
| **37** | Hnrnpm | 0.24 | 3.80E-09 | **37** | Ech1 | -0.20 | 9.66E-18 |
| **38** | Serpine2 | 0.23 | 2.21E-11 | **38** | Mpz | -0.20 | 1.55E-05 |
| **39** | Ednrb | 0.23 | 9.28E-09 | **39** | Sema3c | -0.20 | 3.77E-17 |
| **40** | Cdkn1c | 0.23 | 6.40E-08 | **40** | Olfml2a | -0.20 | 1.28E-14 |
| **41** | S100a11 | 0.23 | 1.68E-09 | **41** | Pmp22 | -0.19 | 1.18E-10 |
| **42** | Mif | 0.22 | 1.15E-08 | **42** | Mbp | -0.19 | 3.82E-09 |
| **43** | H2afy | 0.22 | 3.65E-12 | **43** | Chmp2a | -0.19 | 3.35E-13 |
| **44** | Dstn | 0.22 | 7.08E-14 | **44** | Cltb | -0.18 | 1.17E-15 |
| **45** | Pdap1 | 0.22 | 3.23E-09 | **45** | Mal | -0.18 | 4.35E-20 |
| **46** | Sox4 | 0.22 | 2.97E-11 | **46** | Rpl23a-ps3 | -0.18 | 6.84E-13 |
| **47** | Ran | 0.22 | 2.01E-06 | **47** | Wdr89 | -0.18 | 3.58E-13 |
| **48** | Pkm | 0.22 | 5.18E-11 | **48** | Gm8730 | -0.17 | 7.23E-16 |
| **49** | Col14a1 | 0.22 | 4.07E-12 | **49** | Cisd3 | -0.17 | 9.79E-12 |
| **50** | Nap1l1 | 0.22 | 6.57E-10 | **50** | Rarres2 | -0.17 | 9.54E-11 |

#### Supplementary Table 8. Common HIRA KO *vs* HIRA WT differentially expressed (DE) genes between melanoblast and glial cells from top 50 genes

| Common upregulated genes | Encoded protein | Common downregulated genes | Encoded protein |
|---|---|---|---|
| Fabp5 | Fatty acid binding protein 5 (epidermal) | Fabp7 | Fatty acid binding protein 7 |
| AY036118 | | Gm11808 | |
| Pkm | Pyruvate kinase; muscle | Crip2 | Cystein rich protein 2 |
| Gm42418 | lnRNA gene | Rps27rt | Ribosomal protein S27 |
| Tubb3 | Tubulin, beta 3 Class III | Gm2000 | |
| H2afz | H2A.Z variant histone | Rpl36 | Ribosomal protein L36 |
| H1f0 | H1.0 linker histone | Rpl10 | Ribosomal protein L10 |
| Marcksl1 | Macrophase myristoylated alanin0rich C kinase like 1 | Gm8730 | |
| Mif | Macrophage migration inhibitory factor | Cox20 | Cytochrome C oxidase assembly factor COX20 |
| Hmgb2 | High mobility group box 2 | Chmp2a | Charged multivesicular body protein 2A |
| Gnas | Guanine nucleotide binding protein alpha stimulating | Wdr89 | WD repeat domain 89 |
| Sub1 | Activated RNA polymerase II transcriptional coactivator p15 | Fau | FAU Ubiquitin like and ribosomal protein S30 fusion |
| Nap1l1 | Nucleosome assembly protein 1-like 1 | Rpl23a-ps3 | Ribosomal protein 23a |
| Cbx3 | Chromobox 3 / Heterochromatin protein 1 gamma (HP1g) | Rplp1 | Ribosomal protein lateral stalk subunit P1 |
| Mdk | Midkine (Neurite growth-promoting factor 2) | Ndufb10 | NADH:Ubiquinone oxidoreductase subunit B10 |
| Ran | Ras-related nuclear protein (GTP-binding) | Lmna | Lamin A/C |
| | | Hypk | Huntingtin interacting protein K |
| | | 2410015M20Rik | |

**Supplementary Table 9. Primary antibodies used in IHC**

| Primary antibody | Species | Supplier | Dilution factor |
|---|---|---|---|
| BrdU | Rat | Abcam ab6326 | 1/500 |
| H3 C-terminal | Rabbit | Active Motif 39163 | 1/5000 |
| Histone H3.3 | Rabbit | Millipore 09-838 | 1/1000 |
| RFP (tdTomato) | Rabbit | Tebu-bio 600-401-379 | 1/100 |
| Sox10 | Rabbit | Abcam ab155279 | 1/1000 |
| TRP2 / DCT (D18) | Goat | Santa Cruz sc-10451 | 1/200 |

**Supplementary Table 10. Secondary antibodies used in IHC**

| Secondary antibody and Species | Supplier |
|---|---|
| AF 488 Donkey anti-Goat IgG H+L | Invitrogen A11055 |
| AF 594 Donkey anti-Mouse IgG H+L | Life Technologies A21203 |
| AF 594 Donkey anti-Rat IgG H+L | Life Technologies A21209 |
| AF 594 Donkey anti-Rabbit IgG H+L | Life Technologies A21207 |
| AF 488 Donkey anti-Sheep IgG H+L | Invitrogen A11015 |

**Supplementary Table 11. Technical information on samples used in scRNA seq**

| Experiment | Sample | Sex | Estimated number of input cells | Number of cells analysed in Cell Ranger | Sequencing facility | Illumina HiSeq4000 sequencing parameters* |
|---|---|---|---|---|---|---|
| Chromium 1 | WT1 | F | 7000 | 1549 | Edinburgh Genomics | 75PE, 290M reads per lane |
| | KO1 | M | | 838 | | |
| Chromium 2 | WT2 | M | 10000 | 3639 | UCSD IGM Genomics Center | 26x8x98, 325M reads per lane |
| | WT3 | F | | 3523 | | |
| | KO2 | F | | 4310 | | |
| | KO3 | F | | 4674 | | |

**Supplementary Table 12. Primary antibodies used in western blotting**

| Primary antibody | Species | Supplier | Dilution factor |
|---|---|---|---|
| Acetyl-a-Tubulin (Lys40) | Rabbit | Cell Signaling Technology 5335 | 1/1000 |
| ASF1a | Rabbit | Cell Signaling Technology 2990s | 1/500 |
| BIII tubulin (Tuj1) | Mouse | Promega G7121 | 1/1000 |
| CAIN (Cabin1) | Rabbit | Abcam ab3349 | 1/1000 |
| c-Kit | Goat | R&D systems AF1356 | 1/400 |
| GAPDH | Mouse | Abcam ab9484 | 1/1000 |
| HIRA (WC119) 2mg/ml | Mouse | In house, Hall *et al.** | 1/1000 |
| Lamin A/C | Rabbit | Cell Signaling Technology 2032S | 1/1000 |
| MITF | Mouse | Abcam ab12039 | 1/1000 |
| TRP2 (DCT) | Goat | Santa Cruz sc-10451 | 1/200 |
| a-Tubulin | Mouse | Sigma T9026 | 1/10000 |
| Sox10 | Rabbit | Abcam ab155279 | 1/1000 |
| PCNA | Rabbit | Santa Cruz sc-7907 | 1/200 |
| Actin | Mouse | Sigma-Aldrich A1978 | 1/50000 |

*Hall, C., et al. 2001. HIRA, the human homologue of yeast Hir1p and Hir2p, is a novel cyclin-cdk2 substrate whose expression blocks S-phase progression. Mol. Cell. Biol. 21:1854–1865.

**Supplementary Table 13. Secondary antibodies used in Western blotting.**

| Secondary antibody and Species | Supplier |
|---|---|
| Donkey anti-goat IgG-HRP | Santa Cruz Biotech sc-2020 |
| Goat anti-mouse IgG, HRP linked | Dako P0447 |
| Goat anti-rabbit IgG, HRP linked | Cell Signaling Technology 31460 |
